# Shrew evolution in hybrid zones: a meiotic undecavalent as an exclusive chromosomal supermultivalent and its weak effect on fertility

**DOI:** 10.1101/2023.07.24.550268

**Authors:** Sergey Matveevsky, Oxana Kolomiets, Nikolay A. Shchipanov, Svetlana V. Pavlova

**Affiliations:** Cytogenetics Laboratory, Vavilov Institute of General Genetics, Russian Academy of Sciences, Gubkin str. 3, Moscow 119991, Russia; Laboratory of Population Ecology, A.N. Severtsov Institute of Ecology and Evolution, Russian Academy of Sciences, Leninskiy pr. 33, Moscow 119071, Russia

**Author notes:** Address correspondence to Sergey Matveevsky and Svetlana Pavlova. (SM), (SP).

**Keywords:** Sorex, multivalent, meiosis, synaptonemal complex, nuclear architecture

## Abstract

Hybrid zones are “natural laboratories” for studying speciation. In the common shrew *Sorex araneus*, the hybrid zone between extremely divergent in karyotypes the Moscow and Seliger chromosomal races is unique, because complex heterozygotes (interracial hybrids) form the longest meiotic configuration consisting of 11 chromosomes with monobrachial homology (undecavalent or chain-of-eleven – CXI). Different studies suggested that such a multivalent would negatively affect meiotic progression and, in general, would significantly reduce fertility. In this work, using immunocytochemical and electron microscopy methods, we investigated chromosome synapsis, recombination, and meiotic silencing in pachytene spermatocytes carrying undecavalent. Despite some abnormalities detected in spermatocytes, such as single associations of chromosomes, stretched centromeres in the multivalent, shifted recombinational peak towards distal parts of chromosomal arms of superchain, heterozygous shrews were able to form a large number of morphologically normal and active spermatozoa. Possible low stringency of pachytene checkpoints, proper segregation of homologous chromosomes, and the ability of hybrids to form mature germ cells imply rapid evolutionary fixation and circulation of Rb chromosomes within shrew populations, leading to a variety of chromosomal races.

## 1 INTRODUCTION

Chromosomal rearrangements are the most important subject for evolutionary research because they lead to changes in chromosome structure, gene arrangement, and therefore differentiation of karyotypes (King, 1993 and refs therein). Experimental or natural hybrid individuals are a good model for evaluation of their viability and morphological, physiological, biochemical, genetic, meiotic, and other features in comparison with parental species/karyoforms and thereby determine the degree of reproductive isolation between divergent taxa. According to the concept of chromosomal speciation (Vorontsov, 1960; White, 1978, 1982; King, 1993 and others), chromosomally heterozygous organisms are reproductively inferior to parental forms as a result of defects in gametogenesis, including meiotic failures (reviews: Dobigny et al., 2015; Bakloushinskaya, 2016; Pavlova & Searle 2018). One of convenient models for investigating the consequences of reproductive isolation is heterozygotes (hybrids) carrying a single or multiple Robertsonian (Rb) translocations in karyotypes (reviewed by Matveevsky & Kolomiets 2020). Rb translocations, which were first described by Robertson in 1916, are centric fusion of two single-armed chromosomes (acrocentrics) resulting in a “de novo” biarmed chromosome (metacentric) or a opposite process, centric fission, when two single-armed chromosomes arise from a biarmed one. Rb translocations are the most common type of structural chromosomal rearrangements that lead to changes of 2*n* in karyotypes. In heterozygous individuals, the arms of *de novo* biarmed chromosomes find their homologs in both single-and biarmed chromosomes in the first meiotic prophase, thereby forming a multivalent configuration. The number of multivalents, just as the number of elements in a multivalent, can affect the fertility of heterozygotes to various degrees (Kolomiets et al., 1985, 2023; Bogdanov et al., 1986; Ratomponirina et al., 1988; Lyapunova et al., 1990; Said et al., 1993; Johannisson, Winking, 1994, Sharma et al., 2003). Karyotypic polymorphism of the Rb type is widespread among animals including mammals: *Mus musculus domesticus* (Gropp et al., 1972; Gropp & Winking 1981; Pialek et al., 2005), *Ellobius tancrei* and *E. alaicus* (Romanenko et al., 2019; Bakloushinskaya et al., 2019), *Eulemur* species (Djlelati et al., 1997; Rumpler, 2004), and the subject of the present study: the common shrew *Sorex araneus* Linnaeus, 1758 (Searle & Wójcik, 1998).

The latter represents one of the well-documented examples of exceptional Rb variation among mammals (Searle 1993; Searle et al., 2019). Within its extensive Euro-Asiatic range—from British Isles and France up to the southwestern part of Yakutia (Eastern Siberia, Russia)—the species is subdivided into numerous parapatric groups of local populations, with each group having a distinct karyotype: so-called chromosomal races (Searle, 1993; Mishta & Searle, 2019). The diploid number of chromosomes, 2*n*, of the species varies from 20 to 33 due to centric fusions and/or fissions (Rb) of 10 single-armed autosomes (*g, h, i, k, m, n, o, p, q,* and *r*), constituting the variable part of a karyotype (Figure 1A–C). Four pairs of biarmed autosomes—*af, bc, jl,* and *tu*—and the sex chromosomes (XX in females and XY1Y2 in males) do not vary (Searle et al., 1991).

**FIGURE 1.**
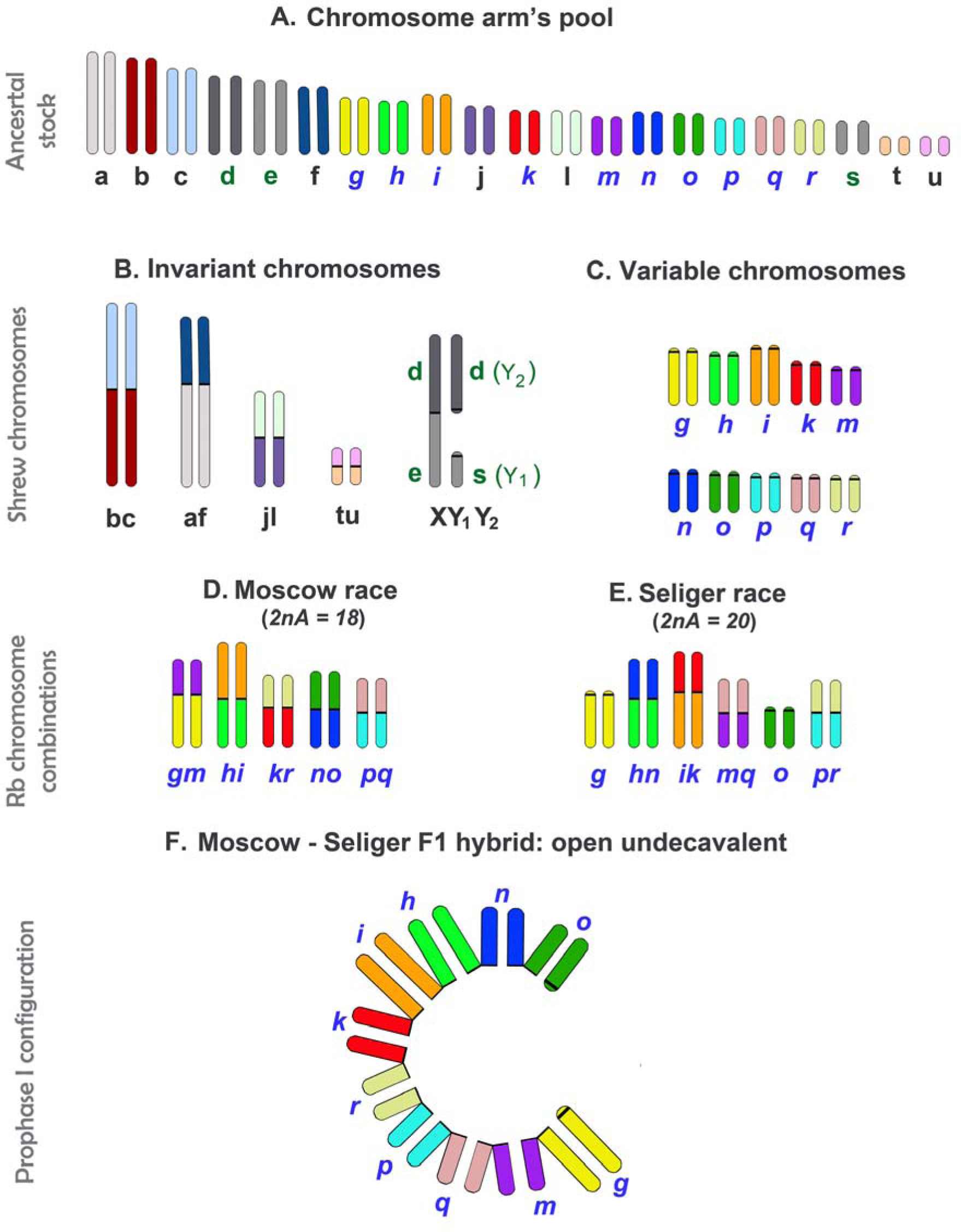
Common-shrew chromosomes. Schemes of ancestral chromosome arms (A) and invariant (B) and variable (C) parts of a karyotype; karyotypic variants of the Moscow (D) and Seliger (E) chromosomal races and the meiotic configuration (undecavalent) in an F1 interracial hybrid (F). The labeling of chromosome arms with letters is given in accordance with the nomenclature of the standard *S. araneus* karyotype (Searle et al., 1991).

The races differ from each other by race-specific chromosome arm combinations resulting from Rb translocations or/and WART (whole-arm reciprocal translocation between two metacentrics); 76 distinct chromosomal races are recognized now (Bulatova et al., 2019). In an area of contact between racial geographic ranges, shrews can meet and produce natural interracial hybrids (heterozygotes); to date, 36 chromosomal hybrid zones of the common shrew have been identified (Fedyk et al., 2019). Such zones are “natural laboratories” for studying microevolutionary processes of early stages of species differentiation (Hewitt, 1988); moreover, chromosomal hybrid zones in *S. araneus* are “tension zones,” which are maintained by a balance between dispersal and selection against hybrids (Searle & Wójcik, 1998). In the hybrid zones, natural “simple” and “complex” heterozygotes (hybrids) have been found (Wallace & Searle, 1990; Fedyk et al., 2019). The former produce one or more trivalents at meiotic prophase I, whereas complex heterozygotes form chain or ring configurations (multivalents) consisting of four or more elements.

Heterozygous shrews from the karyotypically similar Moscow–West Dvina hybrid zone theoretically should form an open CXI of the same length: *gm/mq/**qr**/pr/**ip**/ik/**hk**/hn/no/o* (see D section in Figure 2); nonetheless, neither an F1 hybrid karyotype nor any meiotic data have been presented so far (Orlov et al., 2013). The hybrid zone between the Moscow and Seliger chromosomal races (Central European Russia) is of great interest because these races are extremely divergent in the karyotype (Bulatova et al., 2000 2007; Pavlova et al., 2008). They differ in nine monobrachial metacentrics and two acrocentrics (the Moscow race: ***gm, hi, kr, no, pq****, jl*, 2na=18, and the Seliger race: ***g, hn, ik, mq, o, pr****, jl,* 2na*=*20) (Figure 1D,E) such that their F1 hybrids should produce extremely long chain-of-eleven (CXI) configuration *gm/mq/pq/pr/kr/ik/hi/hn/no/o*, an open undecavalent, in meiosis I (Figure 1F).

**FIGURE 2.**
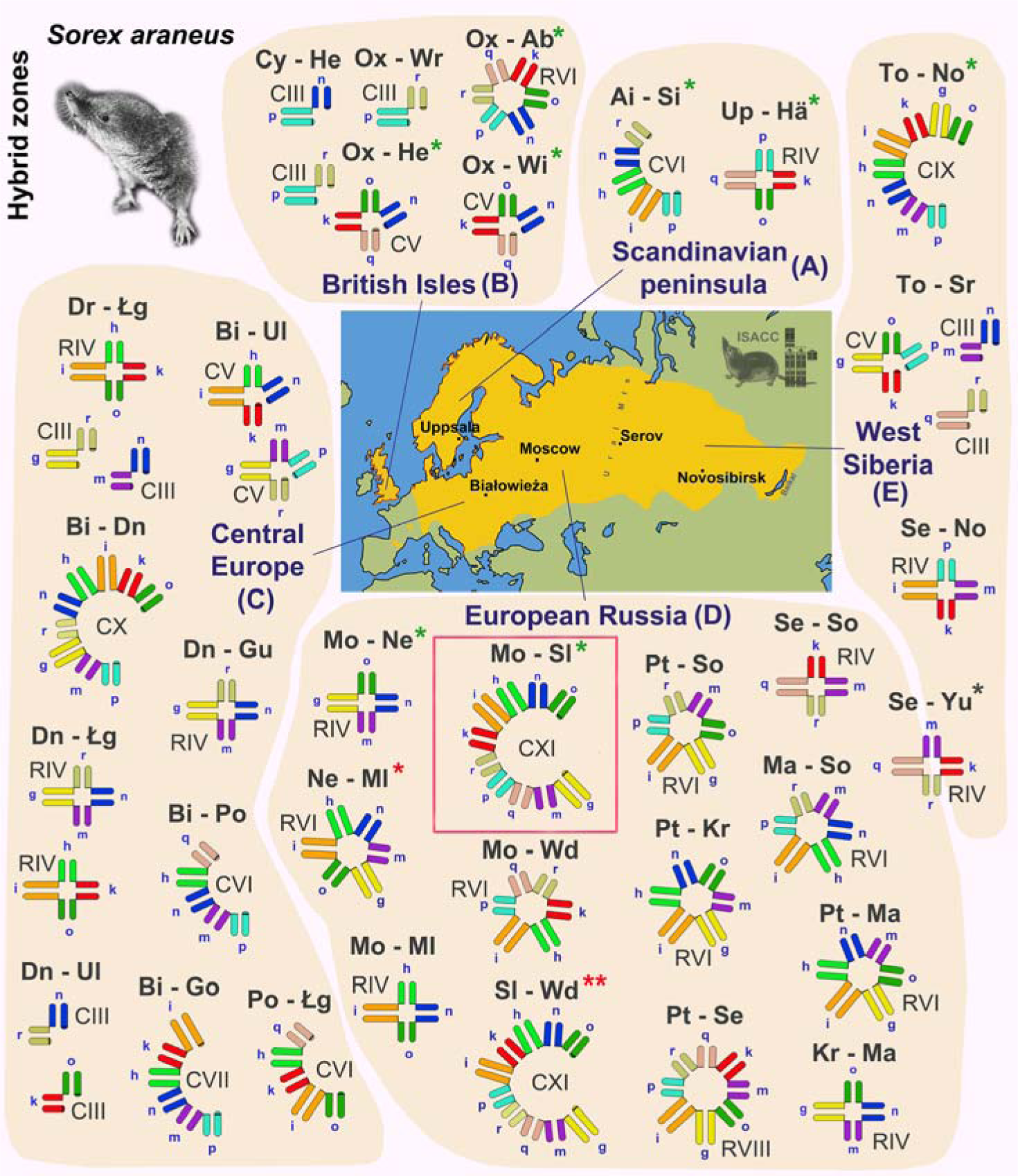
Meiotic configurations at prophase I of heterozygous shrews from hybrid zones. The color of the arms corresponds to the chromosomes in Figure 1. Expected meiotic configurations based on known karyotype data (Fedyk et al., 2019). Undecavalent examined in this work is framed in red. The green asterisks indicate hybrid zones where synaptonemal complexes were studied. A hybrid zone where no hybrid individuals were found marked by single red asterisk. Double red asterisks indicate a hybrid zone, where no primary data demonstrated (no karyotype, no results of meiotic studies). A gray asterisk indicates a race distributed in two geographic regions. Meiotic configurations: C – chain, R – ring; for example, CXI – chain-of-eleven or RVII – ring-of-seven. <u>Hybrid zones and types of</u> <u>multivalents.</u> **A. Scandinavian Peninsula. (1)** Abisko (Ai) – Sidensjö (Si): open hexsavalent (CVI). **(2)** Uppsala (Up) – Hällefors (Hä): closed tetravalent (RIV). **B. British Isles. (3)** Oxford (Ox) – Wirral (Wi): open pentavalent (CV). **(4)** Oxford (Ox) – Hermitage (He): open pentavalent (CV) and trivalent (CIII). **(5)** Oxford (Ox) – Wrentham (Wr): trivalent (CIII). **(6)** Chysauster (Cy) – Hermitage (He): trivalent (CIII). **(7)** Oxford (Ox) – Aberdeen (Ab): closed hexsavalent (RVI). **C. Central Europe. (8)** Drużno (Dr) – Łęgucki Młyn (Łg): closed tetravalent (RIV) and two trivalents (CIII). **(9)** Białowieża (Bi) – Ulm (Ul): two open pentavalents (CV). **(10)** Białowieża (Bi) – Drnholec (Dn): open decavalent (CX). **(11)** Drnholec (Dn) – Guzowy Młyn (Gu): closed tetravalent (RIV). **(12)** Drnholec (Dn) – Łęgucki Młyn (Łg): two closed tetravalents (RIV). **(13)** Drnholec (Dn) – Ulm (Ul): two trivalents (CIII). **(14)** Białowieża (Bi) – Popielno (Po): open hexsavalent (CVI). **(15)** Białowieża (Bi) – Gołdap (Go): open heptavalent (CVII). **(16)** Popielno (Po) – Łęgucki Młyn (Łg): open hexsavalent (CVI). **D. European Russia. (17).** Moscow (Mo) – Neroosa (Ne): closed tetravalent (RIV). **(18).** Moscow (Mo) – Seliger (Sl): open undecavalent (CXI). **(19).** Neroosa (Ne) – Mologa (Ml): closed hexsavalent (RVI). **(20).** Moscow (Mo) – Mologa (Ml): closed tetravalent (RIV). **(21).** Moscow (Mo) – West Dvina (Wd): closed hexsavalent (RVI). **(22).** Seliger (Sl) – West Dvina (Wd): open undecavalent (CXI). **(23).** Petchora (Pt) – Sok (So): closed hexsavalent (RVI). **(24).** Petchora (Pt) – Kirillov (Kr): closed hexsavalent (RVI). **(25).** Petchora (Pt) – Serov (Se): closed oktavalent (RVIII). **(26).** Serov (Se) – Sok (So): closed tetravalent (RIV). **(27).** Manturovo (Ma) – Sok (So): closed hexsavalent (RVI). **(28).** Petchora (Pt) – Manturovo (Ma): closed hexsavalent (RVI). **(29).** Kirillov (Kr) – Manturovo (Ma): closed tetravalent (RIV). **(30).** Serov (Se) – Yuryuzan (Yu): closed tetravalent (RIV). **E. West Siberia. (31).** Tomsk (To) – Novosibirsk (No): open nonavalent (CIX). **(32).** Tomsk (To) – Strelka (Sr): open pentavalent (CV) and two trivalents (CIII). **(33).** Serov (Se) – Novosibirsk (No): closed tetravalent (RIV).

Previously, a CXI meiotic configuration has been detected in the Moscow–Seliger hybrid male by light and electron microscopy (Pavlova et al., 2008). This complex heterozygote was unusual, because carried an additional WART rearrangement, and thus the chain-of-eleven configuration *gm/**hm**/hi/ik/kr/pr/pq/**nq**/no/o* differed from the expected one for F1 interracial hybrids. Therefore, at the moment, the open undecavalent or CXI in Moscow–Seliger F1 hybrids found in natural populations, represents an exclusive meiotic configuration or a supermultivalent with the largest number of elements (Figure 2).

The use of immunocytochemical tools to examine multivalents in prophase I of meiosis makes it possible to reveal patterns of chromosome synapsis as well as organization of centromeres, recombination features, the distribution of epigenetic markers, structural features of chromatin, and nuclear architectonics (Manterola et al., 2009; Matveevsky et al., 2015; Berrios, 2017; Ribagorda et al., 2019). Until now, to our knowledge, immunocytochemical data on meiotic divisions of heterozygous shrew males have been obtained only from two hybrid zones out of 36 known: Novosibirsk–Tomsk and Moscow–Neroosa (Karamysheva et al., 2007; Matveevsky et al., 2012; Belonogova et al., 2017). Consequently, new meiotic data from hybrid zones having different karyotypic complexity of parental races are important. The meiotic undecavalent of F1 hybrids from the Moscow–Seliger hybrid zone is an excellent model for immunocytochemical studies and evaluation of possible effects of chromosomal heterozygosity and multiple chromosomal rearrangements on fertility parameters.

In this work, for the first time, we present data from a pachytene analysis of extremely long meiotic configurations, i.e., undecavalents, while focusing on chromosomal synapsis, meiotic silencing, and meiotic recombination and describe structural features of spermatozoa in complex heterozygotes (F1 hybrids) between the Moscow and Seliger chromosomal races of the common shrew *S. araneus*.

## 2 MATERIAL AND METHODS

### 2.1 Animals

Two examined complex heterozygous males of the common shrew *Sorex araneus* were collected in April 2015 in an area of previously described hybrid zone between the Moscow and Seliger chromosomal races (Bulatova et al., 2011) in the Tver region of Central part of European Russia. One homozygous shrew male (Moscow race) was used to compare some meiotic parameters.

### 2.2 Metaphase chromosomes

Suspensions of metaphase chromosomes were obtained from primary fibroblast cell cultures that were derived from tail biopsies of animals and established following the standard cell culture protocol (Freshney, 2010) with some modifications described earlier (Pavlova et al., 2018). Standard trypsin-Giemsa staining technique (GTG-banding) was used to identify each chromosome arm according to the standard nomenclature for the *S. araneus* karyotype (Searle et al., 1991).

### 2.3 Synaptonemal complexes preparing

Synaptonemal complexes (SCs) were obtained according to Navarro with colleagues (1981) with some modifications (Kolomiets et al., 2010). Testes from two Moscow-Seliger hybrid males were processed. The tunic, fat and large vessels were removed and in Eagle’s medium without glutamine the seminiferous tubules were disaggregated using tweezers. Then the cell suspension homogenized with automatic pipette and was transferred to a centrifuge tube and the volume of the suspension was adjusted to 10 ml. After centrifugation of the suspension at 1500 rpm the supernatant was discarded, the precipitate was diluted with Eagle’s medium to a volume of 3 ml, and the suspension was homogenized. Drops of a hypotonic solution of sucrose (0.1M) were applied to poly-L-lysine coated or plastic coated slides. Then the cell suspension was dropped onto these sucrose drops using automatic pipette. After 3-4 minutes, the drops were combined and then evenly distributed over the entire surface of the slide.

Then slides were transferred to the surface of a cooled plate and dried under a cold fan. Next, the cell suspension in the slide was fixed with ice-cold 4% paraformaldehyde (4%, pH9.2). The slides were washed in a 0.4% Photoflo solution three times for 1-2 minutes and dried in air.

### 2.4 Antibodies and Immunostaining

Slides were rinsed one or two times for 2-3 min in phosphate buffer saline (PBS) and incubated overnight at 4°C with the primary antibodies diluted in antibody dilution buffer (3% bovine serum albumin - BSA, 0.05% Triton X-100 in PBS): mouse anti-MLH1 (mutL homolog 1; 1:50, Abcam, Cambridge, UK), rabbit polyclonal anti-SYCP1 (1:400, Abcam), rabbit polyclonal anti-SYCP3 (1:400, Abcam, Cambridge, UK), human anticentromere antibody CREST (Calcinosis Raynaud’s phenomenon, Esophageal dysmotility, Sclerodactyly, and Telangiectasia) (1:500, Fitzgerald Industries International, Acton, MA, USA), and mouse anti-phospho-histone H2AX (also known as γH2AFX; serine 139 -phosphorylated gamma histone H2A.X) (1:1000, Abcam, Cambridge, UK). After washing in PBS, we used the following corresponding secondary antibodies diluted in PBS: FITC-conjugated bovine anti-rabbit IgG (1:500, Santa Cruz Biotechnology, Santa Cruz, CA, USA), goat anti-rabbit Alexa Fluor 488 (1:400, Invitrogen Corporation, Carlsbad, CA, USA), goat anti-human Alexa Fluor 546 (1:400, Invitrogen Corporation) and goat anti-mouse Alexa Fluor 555 (1:200, 1:1000, Invitrogen Corporation). Then slides were PBS and immersed into Vectashield with 4′,6-diamidino-2-phenylindole (DAPI) (Vector Laboratories, Burlingame, CA, USA). Slides were analyzed using a fluorescence light microscope Axio Imager D1 (Carl Zeiss, Jena, Germany). Single-round and multi-round immunostaining procedure was described previously (Matveevsky et al., 2016, 2021).

### 2.5 Silver nitrate staining and electron mycroscopy

The plastic (Falcon film) coated slides were stained with 50% AgNO3 solution in a humid chamber at 56 °C for 2-3 hours. The slides were washed in distilled water three times for 5-10 minutes and then air-dried. The stained slides were observed in a light microscope Axioskop 40 (Carl Zeiss, Jena, Germany), suitably spread cells were selected, and plastic circles were cut out with a diamond tap and transferred onto grids. The slides were examined under JEM 1011 electron microscope (JEOL, Tokyo, Japan).

### 2.6 Diakinesis

Meiotic chromosome preparations for late prophase I (diakinesis) – metaphase II were made according to a centrifuge free procedure adapted from Williams et al. (1971).

### 2.7 Spermatozoa analysis

Chromosomal homozygotes and heterozygotes of shrews were used for an analysis of the structure, motility, and quantity of spermatozoa. To examine the structure of spermatozoa, we employed the same slides on which we obtained the spreads. The analysis was carried out by electron microscopic and immunocytochemical (nonspecific staining) methods.

For the assay of sperm motility, the material was taken from the epididymis that was ground in PBS in a Petri dish at room temperature. The suspension of spermatogenic cells was placed under a coverslip. Video sequence acquisition was performed by means of a Canon PowerShot A640 digital camera (Canon, Tokyo, Japan) under an Axioskop 40 microscope (Carl Zeiss, Jena, Germany). Sperm motility was assessed by subjective analysis of the video-recorded sequences.

Counting of spermatozoa was carried out by determining the number of DAPI-positive spermatozoa heads in micrographs (area of 60 x 100 microns, an objective with a magnification of 100). Because the preparations of spread cells for the homozygotes and heterozygotes were obtained by an identical method, it was possible to compare the results. This subjective type of analysis was performed as an alternative to the method of counting sperm using the Goryaev camera.

### 2.8 Statistical analysis

The statistical analysis of all data was performed using GraphPad Prism 9 software (San Diego, CA, USA). Mean values (M) and standard deviation (SD) were calculated by the descriptive option of the software. P-values were calculated by t-test (for all meiotic recombination data) or Mann–Whitney two-sided non-parametric test (for other parameters).

## 3 RESULTS

### 3.1 Metaphase and synaptonemal complex (SC) karyotypes in shrew complex heterozygotes

Both examined interracial Moscow–Seliger F1 hybrid males had mitotic karyotypes with 2n=22 (the number of chromosome arms or fundamental number NF = 40, and number of autosome arms NF*a* = 36), which consisted of four metacentric pairs of invariant chromosomes (*bc*, *af*, *jl,* and *tu*), sex chromosomes X(*de*) Y1(*s*) Y2(*dv*), and 11 unpaired chromosomes with monobrachial homology *g/gm/mq/pq/pr/kr/ki/hi/hn/no/o* (Figure 3A).

**FIGURE 3.**
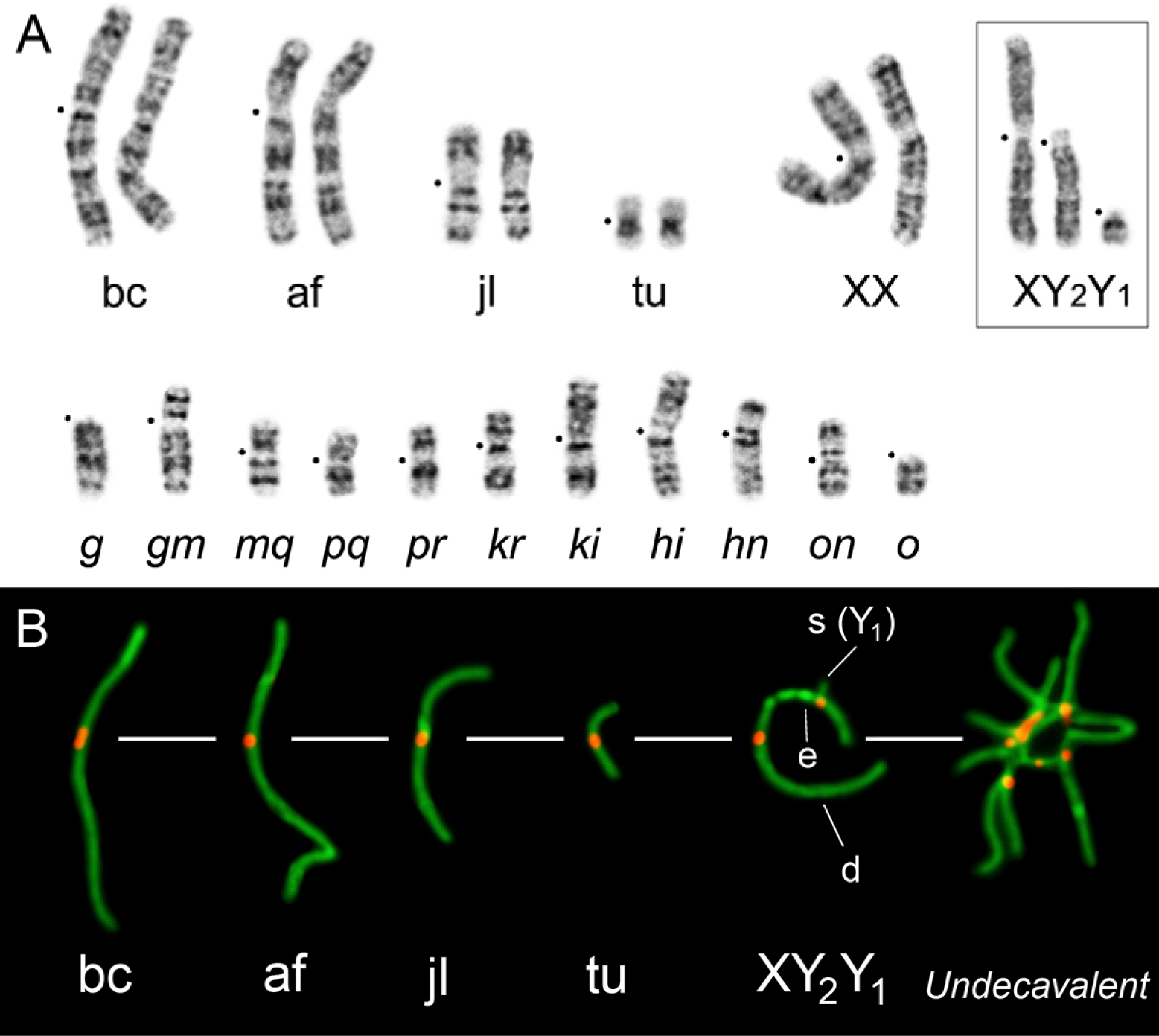
Karyotypes of an F1 hybrid between Moscow and Seliger chromosomal races of the common shrew *S. araneus*. A. The G-banded karyotype with nine monobrachial metacentrics and two acrocentrics forming an undecavalent (chain-of-eleven, CXI) at meiosis. Female sex chromosomes are indicated as XX, and male sex chromosomes XY1Y2 that form a sex trivalent at meiosis are shown in the box. Black dots mark centromere positions. B. The meiotic (SC) karyotype. Chromosomal arms are labeled according to the nomenclature for the standard *S. araneus* karyotype (Searle et al., 1991).

The meiotic or SC karyotype of these hybrids contains four biarmed SC bivalents (*bc*, *af*, *jl,* and *tu*), a sex trivalent (XY1Y2), and a configuration composed of nine biarmed and two single-armed chromosomes, i.e., an undecavalent (chain-of-eleven, CXI): *g/gm/mq/qp/pr/rk/ki/ih/hn/no/o* (Figure 3B).

### 3.2 Chromosome synapsis and irregularities

To determine the structure and behavior of pachytene chromosomes, immunodetection of the SYCP3 protein (a major protein of axial/lateral elements of the SC), SYCP1 (the main component of the transverse filaments in central elements of SCs), and kinetochore proteins (CREST antibody) was performed.

SCs in spermatocytes were analyzed at the pachytene stage. In total, 94 pachytene nuclei were examined, of which 32 were subjected to multiround immunostaining and 12 to silver nitrate staining (for electron microscopy). In the pachytene spermatocyte, four SC bivalents (*bc*, *af*, *jl*, and *tu)*, a sex trivalent, and an open undecavalent with decoded chromosome arms were identified (Figures 4 and 5). None of the four autosome bivalents showed aberrant synaptic patterns.

**FIGURE 4.**
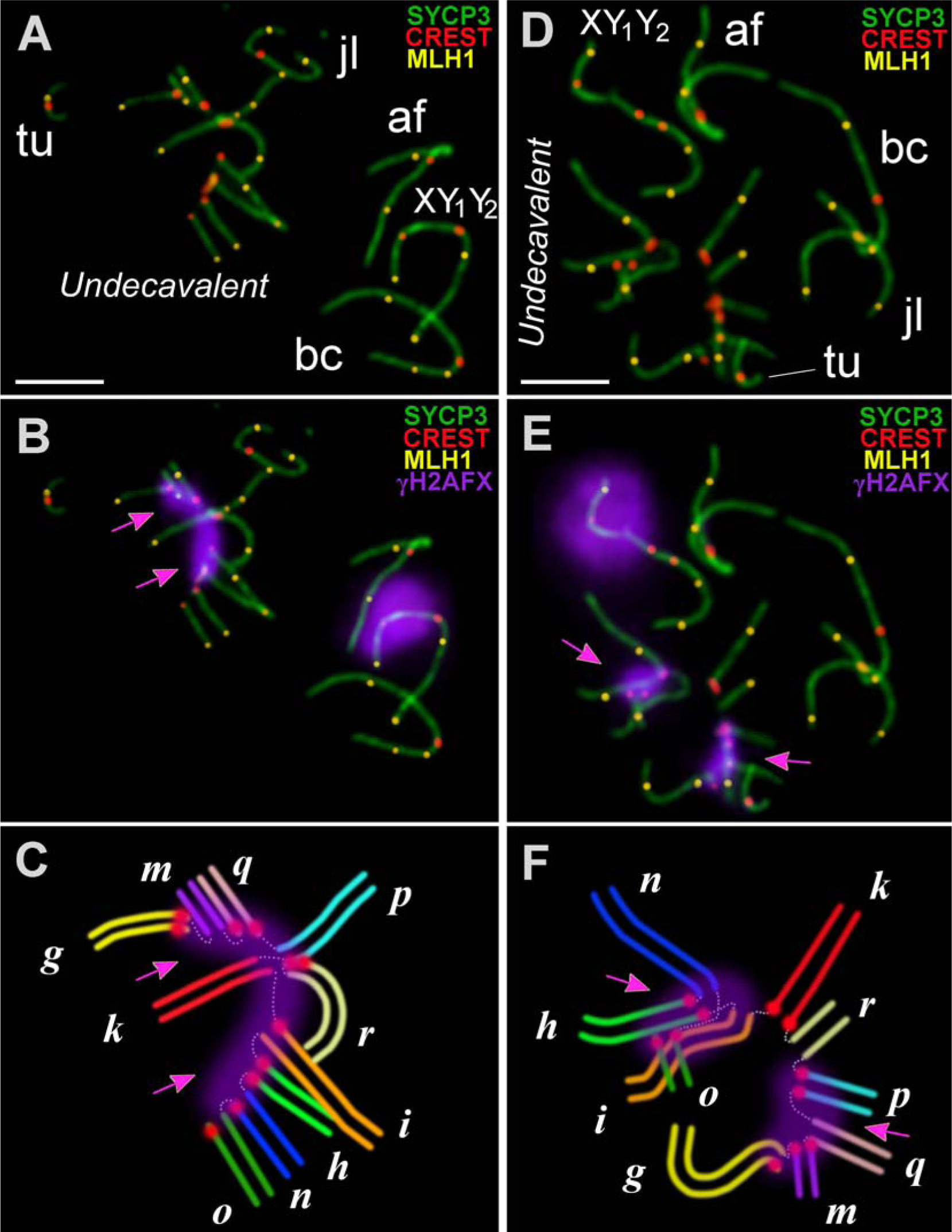
Pachytene spermatocytes (A, B, D, E) and schemes of a chromosomal undecavalent (C, F) in Moscow–Seliger F1 hybrids. SYCP3 (green), CREST (red), MLH1 (yellow), and γH2AFX (violet) are shown. Magenta arrows indicate H2AFX-positive regions of asynapsis within the chromosomal undecavalents (B, C, E, F). SCs are labeled according to chromosomal arms (see Figures 1F and 3). The scheme of the undecavalent (C, F) is based on chromosomes in nuclei (A, B, D, E): each of the arms has a color in accordance with the color of the chromosome arms in Figure 1. Scale bars (A, B, D, E) = 5 μm

**FIGURE 5.**
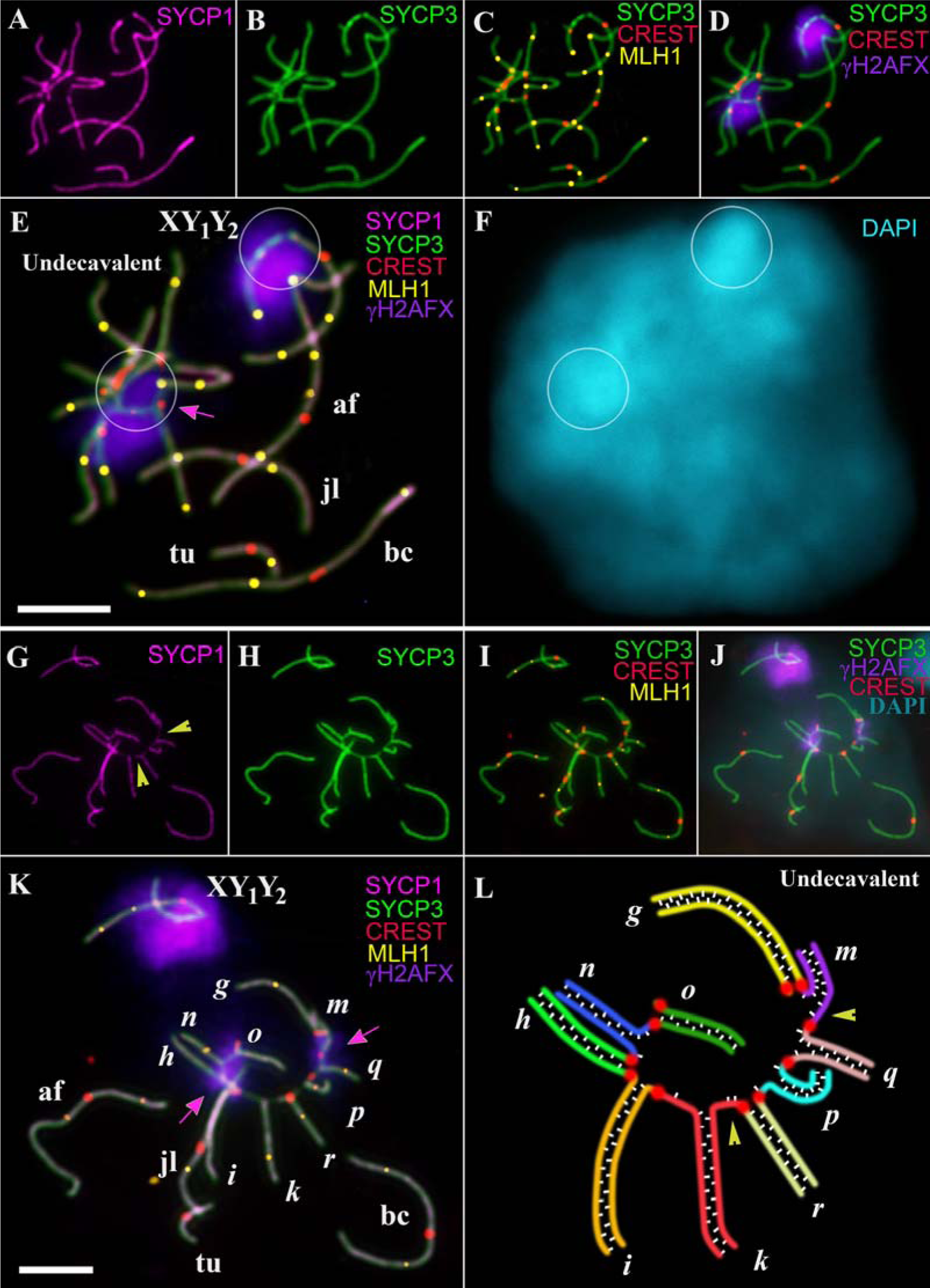
Pachytene chromosomes (nuclei: A–F and G–L), a chromosomal undecavalent (F) in a spermatocyte of a Moscow–Seliger F1 hybrid. SYCP1 (magenta), SYCP3 (green), CREST (red), MLH1 (yellow), γH2AFX (violet), and DAPI (cyan or pale cyan) signals are presented. The same nuclei with different numbers of immunosignals (A–F and G–L) are shown. SCs are labeled according to chromosomal arms (see Figures 1F and 3). White circles denote the same area of the nucleus (E, F) where the DAPI signal is more intense. Magenta arrows indicate H2AFX-positive regions of asynapsis within the chromosomal undecavalents (E, K). The scheme of the undecavalent (L) is based on the meiotic configuration in the nucleus (G–K): each of the arms is colored in accordance with the color of the chromosome arms in Figure 1. SYCP1-positive transverse elements of the central elements of the SCs are indicated by small white sticks (L). SYPC1 is seen in unsynapsed regions (yellow arrowheads) (G, L). Scale bars (E, K) = 5 μm

The sex trivalent (XY1Y2) has structure identical to that described earlier (Pack et al., 1993; Matveevsky et al., 2017). The true (XY1) and translocated (XY2) parts were found to be connected by an asynaptic area (a part of X outside the synaptic regions), which is often thickened and may look like large SYCP3 dots (Figures 3B, 4A,D, 5B,C, and 6B,C). The true part of XY1Y2 gave a stronger DAPI signal (Figure 5E,F). Centromeres of the X and Y2 chromosomes are not aligned in some cases (Figures 4D and S3). The SYCP1 signal was located in the true and translocated parts of XY1Y2 (Figures 5A,B,G,H and 6A,B). The sex trivalent or its true part, as a rule, lies on the periphery of the nucleus (Figures 4A,D and 5C). In 8.6% (n = 76) of nuclei, end-to-end contacts of autosomes with Y1 or the asynaptic part of the sex trivalent were noted (Figures S2 and S3). Some XY1Y2 also contain abnormally long unsynapsed regions (Figures S2 and S3).

An undecavalent was clearly visualized in all spermatocytes. Chromosomal arms in some undecavalents were difficult to identify because the arms overlapped with each other (Figures 3B and S1). In most of the undecavalents, we were able to recognize chromosome arms on the basis of their known length. In the MicroMeasure software, the *g* arm was assigned to the longest of the two acrocentrics involved in this multivalent. Next, the chain was deciphered due to known homologies of the arms (Figures 4A,C,D,F and 5A,L). One-third of the examined undecavalents (30.49%, n = 70) were found to be divided into pieces that lay at a distance from each other (Figures S1, S3, and S4). In the majority of nuclei, the undecavalent was open (chain configuration) (97.42%, n = 70) (Figures 4A,C,D,F and 5K,L). Only two nuclei with undecavalents were observed; there was a ring-like configuration in one case (Figure 5A–E), and in the other case, a part of the undecavalent appeared to be fully synapsed (closed configuration) (Figure S4). Shrews’ chromosomes are quite long, and therefore overlapping of chromosomes was often observed. Except for such cases, nuclei showing true association of an undecavalent with a sex trivalent or autosomes were not found at all.

In undecavalents, centromeres were visualized as dots that lie in the middle of the axial/lateral elements except for two acrocentrics, *g* and *o* (Figures 4A,D, 5I, and S4). In some biarmed chromosomes, the SYCP3 signal was often invisible in unpaired areas of transition from one synapsed arm to another within an undecavalent (Figure 4); this phenomenon can be considered an artifact of the spreading. In 15.9% (n = 77) of undecavalents, the centromeric regions proved to be stretched (an elongate or line-like pattern) (Figure 6). It seemed that parts of the undecavalent lay far apart (a dissociated undecavalent); however, stretched centromeres combined these parts into a single multivalent. It has previously been shown in squashed spermatocytes that such stretched centromeres are not artifacts of the spreading procedure (Gil-Fernández et al., 2020, Matveevsky et al., 2020). A more intense DAPI signal was detected in regions of the undecavalent centromeres (Figure 5E,F).

**FIGURE 6.**
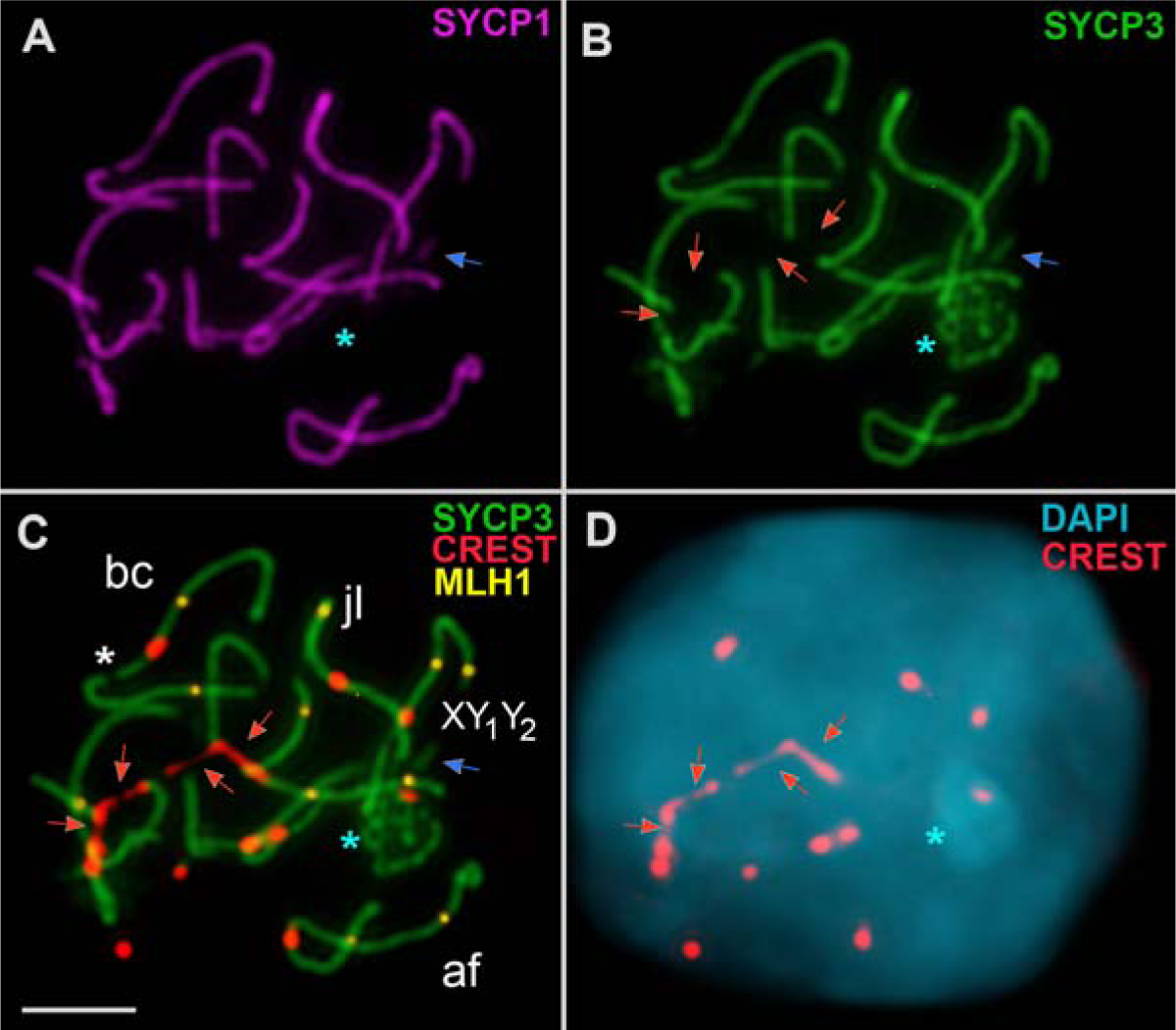
Stretching of centromeric regions in a meiotic undecavalent from a pachytene spermatocyte of a Moscow–Seliger F1 hybrid. SYCP1 (magenta), SYCP3 (green), CREST (red), MLH1 (yellow), and DAPI (cyan) signals are presented. Red arrows indicate stretched centromeric regions within the chromosomal undecavalent. The blue arrow points to the synaptic region of the true part of the sex trivalent. The blue asterisk marks the thickened asynaptic part of the sex trivalent. The asynaptic part of XY1Y2 shows a more intense DAPI signal. The white asterisk marks the gap in the bc SC bivalent. Scale bars = 5 μm

Of note, the SYCP1 protein is localized to some unpaired regions of the undecavalent (Figure 5G,L). In these cases, SYCP1 localization is discontinuous, i.e., SYCP1 is not located along the entire unpaired region. In the only case where a part of an undecavalent was fully synapsed, SYCP1 signal was registered at junctions between the arms, i.e., in those areas where there was asynapsis in a chain configuration (Figure S4B). In these areas, thick SYCP3 axes were observed (Figure S4A), indicating the presence of so-called heterologous (nonhomologous) synapsis. A similar type of heterologous synapsis has been described in a meiotic hexavalent of hybrid mice (Ribagorda et al., 2019).

### 3.3 Meiotic inactivation of unpaired areas of the undecavalents

It is known that chromosomal regions undergo a process of transcriptional inactivation called meiotic silencing of unsynapsed chromatin (MSUC) (Schimenti, 2005; Baarends et al., 2005; Turner et al., 2005). A specific case of MSUC is meiotic inactivation of sex chromosomes (MSCI) (McKee & Handel, 1993; Turner et al., 2002; Hendel, 2004). It has been suggested that in pachytene, inactivation of asynaptic chromosomal regions carrying genes critical for meiotic progression may induce arrest of meiocytes (Turner et al., 2005), whereas wide asynapsis and MSUC may impair MSCI (Mahadevaiah et al., 2008; Burgoyne et al., 2009). To characterize MSUC and MSCI processes in shrew hybrid spermatocytes, we analyzed the localization of the phosphorylated (at serine 139) form of histone H2AX (γH2AFX) and revealed that the γH2AFX signal was absent from autosomes bc, af, jl, and tu at the mid pachytene stage (Figures 4A,B,D,E and 5D,J). Within the sex trivalent, the γH2AFX signal was located in the true part and asynaptic region (Figures 4A,B,D,E and 5D,J), as we described earlier (Matveevsky et al., 2017).

In the undecavalent, as a rule, the γH2AFX signal covered either completely (Figures 4B,C and 5D,E) or partially the asynaptic segments (Figures 4E,F and 5J,K) that formed between the synapsed arms. In some undecavalents, where there was no SYCP3 axis between the synapsed arms, and the γH2AFX signal was absent (Figures 4E,F and 5J,K). This picture can be explained by an artifact of the spreading procedure, or, as previously suggested in a similar case in mice (Ribagorda et al., 2019), by the fact that these regions already experienced the synapsis stage and then prematurely desynapsed, followed by partial elimination of the SYCP3 protein. In the only case where a part of an undecavalent was completely synaptic, there was no γH2AFX signal, thus confirming the absence of asynaptic zones because these regions were in heterologous synapsis (Figure S4A,C). γH2AFX signals in undecavalents and sex trivalents were found to be isolated and did not connect with each other. This pattern was observed in all well-spread nuclei, pointing to topological separation of MSUC and MSCI in hybrid pachytene spermatocytes. This may be functional evidence for the absence of associations between the undecavalent and XY1Y2.

### 3.4 Meiotic recombination

MLH1 foci were used as markers of late recombination nodules and chiasmata (Anderson et al., 1999). It has been proposed that one chiasm per arm is required for successful chromosome segregation (Dumas & Britton-Davidian, 2002). In the haploid set, a Moscow–Seliger F1 hybrid has 20 chromosome arms, and therefore 20 MLH1 foci per one spermatocyte were expected. A deviation from this number was revealed in F1 hybrids, whereas this parameter was close to the expected one in a pure parental individual (Moscow race, NF = 40). The number of MLH1 signals (mean ± SD) per nucleus was 17.52 ± 3.3 (n = 44) for one F1 hybrid and 18.03 ± 3.1 (n = 31) for a second one (no statistically significant difference, p < 0.05), and 20.1 ± 3.5 (n = 66) for a male of the Moscow race. Nevertheless, an individual of the parent race and F1 hybrids differed significantly (p < 0.001). Our results are consistent with the number of recombination nodules per nucleus seen in homozygotes and some heterozygotes from the Tomsk–Novosibirsk hybrid zone (Belonogova et al., 2017).

Long autosomes *bc, af*, and *jl* showed two to four MLH1 foci, while the shortest autosome *tu* had one or no MLH1 foci at all (Figures 4A,D and 5C,I). In the sex trivalent, one MLH1 focus in XY1 and one or two MLH1 foci in XY2 were visible (Figures 4A,D, 5C,I, and S4C). In the undecavalent, 10 expected MLH1 foci should be detectable, but it turned out that on average, there were seven to nine MLH1 foci per multivalent (MLH1 foci were missing on one or two arms) (Figures 4D,E and 5C,E). Nevertheless, in rare cases, an increased number of MLH1 foci was seen in undecavalents owing to additional 2–3 signals in one of the synapsed arms (Figure 4A,B). Of note, those nuclei where stretched centromeres are present in undecavalents contain a reduced number of MLH1 foci (e.g., 12 MLH1 dots per nucleus and four MLH1 dots within the undecavalent in Figure 6C). In the undecavalents, MLH1 foci are mainly situated in the distal or proximal segments and very rarely in the pericentromeric segments.

### 3.5 Diakinesis

Diakinesis/metaphase I spreads of both examined F1 hybrid males had a long chain configuration (CXI) together with four autosomal bivalents *bc, af*, *jl,* and *tu* and the sex trivalent (Figure S5), as expected from the mitotic karyotype. A meiotic chain was observed in all the 30 diakinesis/metaphase I spreads scored, none of which showed autosomal or sex chromosome abnormalities. All the observed configurations are similar to those in heterozygous shrews with the WART rearrangement (Pavlova et al., 2008).

### 3.6 Quantity, morphology, and motility of shrew spermatozoa

A comparison of quantitative and qualitative characteristics of spermatozoa allows us to evaluate potential reproductive success in parental races and hybrids (Swan & Christidis, 1987; Wünsch & Pfennig, 2013; Ishishita et al., 2015; Spangenberg et al., 2017 and others). Given that in Soricidae (Cooper & Bedford, 1976; Green & Dryen, 1976; Mori et al, 1991; Kaneko et al., 2001, 2011; Soon-Jeong et al., 2006; Parapanov et al., 2009), in particular in *S. araneus* (Plöen et al., 1979; Pavlova et al., 2008; Parapanov et al., 2009), spermatozoa have been investigated previously, we were able to compare those data with the results obtained in the current work. For the analysis of spermatozoa, we detected spermatozoa on slides when preparing spread meiotic chromosomes or in a suspension of germ cells obtained from testes.

Unfortunately, we could not count spermatozoa by the standard technique (Goryaev camera); therefore, an alternative method was applied. The number of sperm heads captured by microphotography at ×100 magnification (an approximately 60 x 100 micron area) was calculated for homozygotes and both hybrids. Unexpectedly, the results were almost identical (mean ± SD): 2.97 ± 2.6 (range: 0–12, n = 71) for homozygotes, 2.86 ± 2.3 (range: 0–11, n = 35) for one hybrid, and 2.94 ± 2.9 (range: 0–11, n = 31) for another hybrid. The differences were not statistically significant (P < 0.05).

Shrew spermatozoa proved to have the morphological structure that is typical for mammalian cells (Figure 7A–E for homozygotes and Figure 7F–K for heterozygotes). Any spermatozoon of homozygous and heterozygous individuals has a head and a tail with a middle piece and principal piece (Figure 7A,F). Because spermatozoa were detected on preparations of spread cells, in some spermatozoa, the spreading forces disrupted the membrane in one of the central regions of the tail. In such an area, fibers were visible (Figure 7A,F). Depending on specifics of silver nitrate impregnation during staining, we registered slightly different electron density of stained structures inside the head. Nevertheless, a semitranslucent acrosome and an electron-dense nucleus were clearly visible in each sperm head (Figure 7A–D,F–J). As a rule, the acrosome is heterogeneous, and accordingly the structures of different electron density are visible inside. The base of the head often resembles a structure called the sperm calyx, into which the nucleus is embedded (Figure 7B). In rare cases, we observed "isolated" acrosomes (Figure 7C), which could be interpreted as an artifact of the spreading procedure. Note that the acrosome is not detectable by light microscopy (Figure 7E,K and Pavlova et al 2008); hence, only nuclear parts of the spermatozoa heads were visible. Spermatozoa heads identified by DAPI staining have a variety of shapes, which correspond to different stages of differentiation (Figure 7E,J). The head shapes were identical between the homozygotes and heterozygotes. The heads of shrew spermatozoa are shovel-shaped, and we usually see their frontal plane (Figure 7A–D,F–I). The only case where the sperm head could be visualized in the near-sagittal plane showed an enlarged base of the head and a narrowed acrosome (Figure 7C).

**FIGURE 7.**
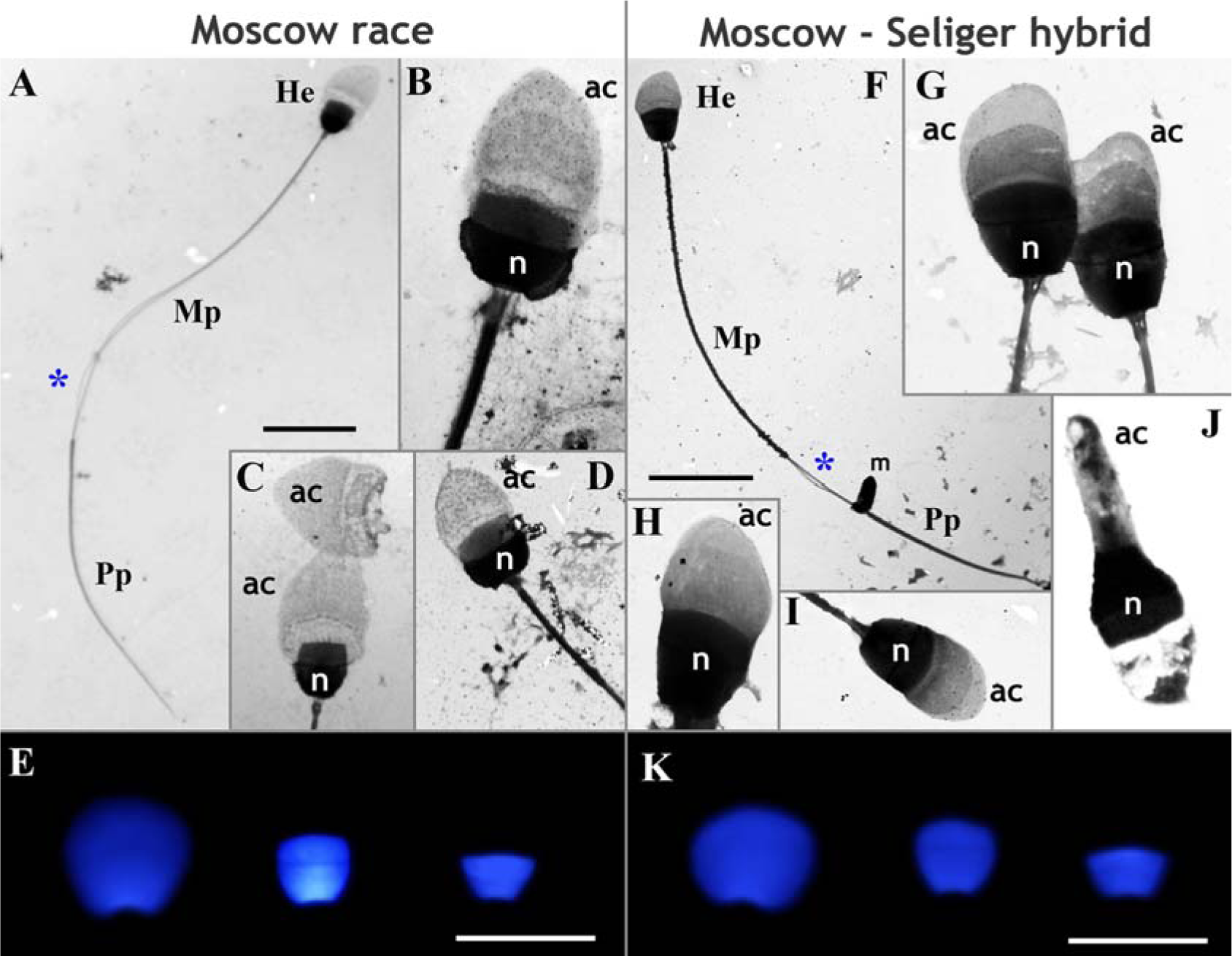
Structure of spermatozoa in the common shrew of the pure Moscow race (A–E) and in Moscow–Seliger F1 hybrids (F–K). Some images were captured after silver nitrate staining (A–D and F–J), and others after DAPI staining (blue) (E, K). Panel J is presented in the near-sagittal plane, and the others in the frontal plane. *Legend.* He: head; n: nucleus, Mp: middle piece, Pp: principal piece, ac: acrosome, m: a part of the burst plasma membrane of the tail. The blue asterisk marks some fibers after the bursting of the plasma membrane of the tail. Explanations are provided in the text. Scale bars (A, F, E, K) = 10 μm

We did not perform quantitative and qualitative analyses of sperm activity, but the presence of motile spermatozoa in homozygous and heterozygous shrews was assessed by means of a video recorded by the digital microscope camera. We identified motile and immobile spermatozoa in both homozygotes (Video S1) and heterozygotes (Video S2). It has been reported that the number of active spermatozoa in homozygous shrews is ∼86.5% (Parapanov et al., 2009), and therefore the presence of immobile spermatozoa can be regarded as a normal pattern. Thus, our data revealed that homozygous and heterozygous shrews have approximately the same numbers of morphologically similar and active spermatozoa, which are probably capable of performing their reproductive function.

## 4 DISCUSSION

We expected that Moscow–Seliger F1 hybrids of the common shrew having a complex multivalent of 11 chromosomes (CXI) would show large-scale meiotic aberrations that activate pachytene arrest or lead to unbalanced segregation and subsequent selection of germ cells. Our results, however, indicated that the picture is different. It is very likely that Moscow–Seliger complex heterozygotes are fertile. The main question is, is there any reduction in fecundity in such complex heterozygotes? We can consider four possible interpretations.

First, synaptic disturbances and chromosome associations can disturb the progression of meiosis (Foreit 1984; Burgoyne & Baker, 1984). Second, pachytene asynapsis (or unsaturated mating) triggers a pachytene checkpoint followed by blockage of meiotic progression (Roeder et al., 2000; Morelli & Cohen 2005; Mahadevaiah et al., 2008; Burgoyne et al., 2009). Third, Rb heterozygotes have an aberrant level of recombination (i.e., the presence/absence of chiasmata) (Davisson & Akeson, 1993; Merico et al., 2003; Dumas et al., 2015), which can lead to unbalanced chromosome segregation and subsequent death of germ cells (King, 1993 and references therein). Fourth, chromosomal multivalents can disturb nuclear architectonics, thereby also possibly leading to abnormal chromosome segregation and cell loss (Berrios et al., 2004; Berrios, 2017). Below we discuss each of these scenarios.

### 4.1 Synapsis and silencing of unpaired regions of chromosomes

Synapsis of chromosomes is an important stage in the progression of meiosis. In particular, this notion has been confirmed on mice carrying mutations in the genes involved in the formation of various SC components. In such mutants, defective synapsis causes pachytene arrest (de Vries et al., 2005; Bolcun-Filas et al., 2007; Hamer et al., 2008).

Some studies on heterozygous shrews from hybrid zones have identified several possible aberrations in the structure and behavior of meiotic chromosomes, including synaptic irregularities:

1. Associations of meiotic chromosomes, including multivalent–XY1Y2, multivalent–autosome, or XY1Y2–autosome;
2. Full asynapsis as univalents in pachytene cells and univalents in diakinesis/MI;
3. Partial asynapsis of meiotic chromosomes, including multivalents, autosomes, or/and XY1Y2.

Association of autosomes and complex SC configurations with sex chromosomes in prophase I of meiosis can reduce fertility or even induce complete sterility (Forejt et al., 1981; Forejt 1984; Burgoyne & Baker, 1984). In the Moscow–Seliger F1 hybrids examined here, we did not detect true associations of the undecavalent with sex trivalent XY1Y2 or autosomes. In our previous study, in Moscow–Neroosa F1 hybrids, a few associations of a tetravalent (ring-of-four, RIV) and a sex trivalent were identified, which resembled an overlap of chromosomes (Matveevsky et al., 2012). In complex heterozygotes obtained in captivity between the Oxford race and Aberdeen race, a multivalent (CVII) was associated with XY1Y2 in four out of 95 spreads (Mercer et al., 1992), whereas in Uppsala–Hällefors hybrids forming the RIV configuration, associations with XY1Y2 and autosomes are observed in 2% and 6% of surface spreads, respectively (Narain & Fredga, 1997). Consequently, it can be concluded that true associations of multivalents with other chromosomes are rare in heterozygous shrews. It has been demonstrated that in some hybrid zones, despite XY1Y2 being associated with autosomes or univalents in different percentages of spreads (Mercer et al., 1992; Fedyk et al., 2005), this phenomenon does not disturb the progression of meiosis. In our work, rare associations of autosomes with the unpaired segment of XY1Y2 in heterozygotes having CXI were also not critical for meiotic progression.

It is thought that monovalence (univalence) may be due to synaptic abnormalities and reduced recombination between completed homologs (Eichenlaub-Ritter, 1996; Ferguson et al., 1996). Indeed, complex Rb heterozygous shrews here shown various levels of univalence frequencies. Nevertheless, univalents, as a rule, have been found at the stage of diakinesis/MI and were rarely seen in pachytene spermatocytes. In particular, no univalents were found at the pachytene stage in case of complex Rb heterozygotes from the Moscow–Seliger (this study) and Moscow–Neroosa (Matveevsky et al., 2012) hybrid zones. In contrast, several cases of the presence of univalents either on autosomes or on sex chromosomes have been observed in pachytene cells of Tomsk–Novosibirsk complex Rb heterozygotes, CIX (Belonogova et al., 2017). In diakinesis/MI nuclei of complex Rb heterozygous, the frequency of univalence varies: for example, 0.14–2.3% and 13.04% in Łęgucki Młyn–Guzowy Młyn hybrids (Banashek et al., 2002) and Drnholec–Białowieża hybrids (Jadwiszczak & Banaszek, 2006), respectively.

In pachytene cells, univalents exemplify whole-chromosome asynapsis; however, partial asynapsis such as unpaired regions of chromosomes can also be identified. γH2AFX is a marker of transcriptionally silenced unsynapsed chromatin regions during meiosis. Previously, this marker was found to be localized in regions of incomplete synapsis for chromosomal translocations (Baarends et al., 2005, Turner et al.,, 2005). In our study, we identified γH2AFX-positive unsynapsed regions in undecavalents. Probably, these silencing regions do not contain critical genes and the meiotic progression of such spermatocytes continues. As noted earlier, the effect of MSUC may be due to both the number and nature of genes that have undergone transcriptional inactivation (Manterola et al., 2009) and this will affect the stringency of the pachytene checkpoint. In the closed tetravalent in Moscow–Neroosa hybrids, areas of slightly elongated centromeres showed no signs of silencing (Matveevsky et al., 2012). Silencing features in shrew undecavalent are very similar to those in mouse hexavalent (Ribagorda et al., 2019).

We found that MSCI and MSUC were not topologically related in the heterozygous nucleus: there was no overlap between γH2AFX clouds belonging to the sex trivalent and the asynapsis of the undecavalent. Considering that aberrant MSUC is capable of violating MSCI (Mahadevaiah et al., 2008; Burgoyne et al., 2009), the facts described probably rule out this possibility in undecavalent-carrying nuclei.

### 4.2 Recombination of meiotic undecavalents likely leads to balanced chromosome segregation

Early notions that unpaired synapsis and aberrant segregation of chromosomes give rise to unbalanced gametes and as a consequence to hybrid sterility and reproductive isolation (White, 1978, 1982; King, 1993) are not always confirmed experimentally (Coyne, 2004). As an alternative, it has been hypothesized that chromosomal rearrangements reduce gene flow by modifying recombination patterns (Rieseber, 2001; Livingstone & Rieseberg, 2004; Borodin et al., 2008, Faria & Navarro, 2010). Recombination is a critical factor in the progression of meiosis, as confirmed by research on mice with mutations in recombination genes. In such mice, defective recombination induces a delay in chromosome synapsis, synaptic disturbances, and subsequent blockage of meiotic progression at zygotene and pachytene stages (Baudat et al., 2000, Romanienko & Camerini-Otero, 2000; Barchi et al., 2005; Kumar et al., 2010; Silva et al., 2013). Several studies show that Rb homozygous mice have a reduced rate of recombination in comparison with all-acrocentric mice (Bidau et al., 2001). Rb heterozygous mice manifest a reduced level of recombination owing to delayed synapsis in trivalents (Davisson & Akeson, 1993; Merico et al., 2003).

A slight decrease in the level of recombination in Moscow–Seliger F1 hybrids of the common shrew may be associated with stretching of the undecavalent (stretched centromeres) in the nucleus; this event could affect the formation of recombination nodules. Nonetheless, in cases of normal centromeres, the number of recombination nodules in multivalents was close to normal (approaching the expected value). Only minimal differences in the number of chiasmata per nucleus have been found between homozygotes and simple and complex heterozygotes in the Drnholec–Białowieża hybrid zone (Jadwiszczak & Banaszek, 2006). In Tomsk–Novosibirsk hybrids, the level of recombination in multivalents was also close to the expected one (Belonogova et al., 2017). In Moscow–Seliger F1 hybrids, the displacement of recombination nodules in the undecavalent was not evaluated in comparison with metacentrics in homozygotes. Nevertheless, in the majority of undecavalents, MLH1 signals were far from the pericentromeric regions and had localization near telomeric sites. These results are consistent with those obtained on Moscow– Neroosa hybrids (our unpublished data) and Tomsk–Novosibirsk hybrids (Belonogova et al., 2017). Those authors suggested that such a recombination pattern in multivalents may be due to the finding that even homozygotes show strong suppression of recombination in proximal regions of chromosomes. That is, this may be some ancestral property of chromosomes. Note that this pattern in multivalents has not always been observed in other mammalian species. For example, in 11% of cases, the localization of recombination nodules is clearly seen at the border of synaptic and asynaptic sites in a hexavalent of heterozygous mice (Ribagorda et al., 2019). A similar distribution of MLH1-positive crossovers is observed in a tetravalent of mole voles (Matveevsky et al., 2015). Distalization of telomeric peaks (shift of MLH1 signals to the distal parts of chromosomes) in the undecavalent may have some evolutionary implications. It is likely that the free acrocentrics’ regions that were once recombination active have become recombination inert within the multivalent. Such sections can be inherited as stand-alone blocks. Thus, these changes may lead to the formation of supergenes or coadaptive gene complexes (Schwander et al., 2014; Thompson & Jiggins, 2014), which enhance the differentiation of isolated animal populations.

It is likely that most undecavalents segregate correctly in anaphase I, although further investigation is needed to estimate the frequency of meiotic nondisjunction in Moscow–Seliger F1 hybrids. We have no data on the fate of undecavalents having elongated centromeres and a reduced level of recombination. It is possible that cells with such multivalents are eliminated, but the large number of spermatozoa indicates likely absence of large-scale cell selection. Therefore, a level of recombination close to normal is probable thus ensuring successful meiotic progression.

### 4.3 Nuclear architecture of hybrid shrew meiocytes: stretched centromeric regions of undecavalents as a possible indicator of a nuclei reorganization

Three-dimensional organization of the genome in cell nuclei may play a key role in the regulation of gene expression and epigenetic processes and may generally implement control over cellular physiological processes (Belyaeva et al., 2017). It is generally accepted that the organization of the nucleus is not random in somatic (Cremer et al., 1993; Parada & Misteli, 2002; Misteli, 2005) or meiotic cells (Berrios & Fernández-Donoso, 1990; Alsheimer et al., 1999; Berrios et al., 2004; Foster et al., 2005; Berrios 2017). Nuclear architecture or the intranuclear landscape characterizes the distribution of chromosome territories within the nucleus (Cremer & Cremer, 2001, 2010; Misteli, 2005; Cremer et al., 2006; Meaburn & Misteli, 2007). Chromosome territories represent spatial domains of different sizes, which specifically occupy a certain volume in the nucleus (Misteli, 2008). The organization of nuclei in meiosis has its own specific features. For example, a meiosis-specific structure, SC, forms a specific chromatin pattern (Hernandez-Hernandez et al., 2009), SCs implement the interaction with the nuclear envelope through shelterin and LINC complexes (Alsheimer et al., 2009; Zetka et al., 2020), and nonsynaptic portions of autosomes and sex chromosomes in prophase I undergo meiosis-specific inactivation processes, MSUC and MSCI (see above first section in Discussion). It can be theorized that nuclear architecture of meiocytes in prophase I is specific for each species (Berrios et al., 2004; Berrios, 2017).

Many studies over the past half century have revealed heterozygote nuclei’s various meiotic configurations (for instance, Kolomiets, et al., 1985, 1986; Ratomponirina et al., 1988; Johannisson & Winking, 1994, 1998; Matveevsky et al., 2015, Ribagorda et al., 2019) in which processes of repair, recombination, and meiotic silencing can be abnormal (Homolka et al., 2007; Mahadevaiah et al., 2008; Burgoyne et al., 2009; Bhattacharyya et al.,, 2013). All this, to one degree or another, could lead to reformatting/deformation of nuclear architecture. In several articles, meiotic nuclei of hybrids have been examined in the context of concepts “nuclear architecture” and “chromosome territories” (Berrios et al., 2014, 2018; Berrios, 2017; Matveevsky et al., 2020). As mentioned above, in 15.9% of undecavalents, at least one centromere looked like a line (stretched centromere) in the present work. This phenomenon could be due to the tension forces that stretch the chromosomes in some undecavalents. The fact is that all chromosomes were found to have points of attachment to the nuclear envelope, and if the distance between the attachment points is greater than the length of metacentric chromosomes in the undecavalent, then centromere stretching is likely to occur. A similar case in intraspecific and interspecific hybrids of mole voles has been documented (Matveevsky et al., 2020); both hybrids had stretched centromeres when located within trivalents, but hyperextension of centromeric regions was observed only in sterile interspecific hybrids. The length of stretched centromeres in the undecavalent of Moscow–Seliger F1 hybrids of the common shrew is comparable to that in trivalents of semifertile intraspecific hybrids of mole voles. It is worth mentioning that in heterozygous shrews from the Tomsk–Novosibirsk hybrid zone, the SC multivalent (CIX) has gaps (Karamysheva et al., 2007; Belonogova et al., 2017). Some of these gaps may have had stretched centromeres. In some shrew heterozygotes, the division of a single meiotic chain into separate multivalents has been noted previously (e.g., in Drnholec–Bialowieza hybrids, Jadwiszczak & Banaszek, 2006), some of which could also be connected by centromeres; thus, these pseudo-autonomous elements could be a part of a single whole. Stretched centromeres can be also considered an indicator of reformatted nuclear architecture of spermatocytes of heterozygous shrews.

Can the altered nuclear architecture affect further progression of meiocytes? Given that most undecavalents here had a single short stretched centromere, it is likely that such spermatocytes did not have problems. The cases where the undecavalent centromeres were slightly longer (for example, in the spermatocyte in Figure 6) implied greater nuclear reformatting. By analogy with mole voles, it is possible that a small percentage of spermatocytes is blocked in their development, but there is no direct evidence. In contrast to interspecific hybrids of mole voles, which do not have spermatozoa at all and are sterile, intraspecific mole vole hybrids have spermatozoa and low fertility (Matveevsky et al., 2020). The latter case is similar to what we observed in heterozygous shrews.

Thus, the observed stretching of centromeres in heterozygous shrews can be considered a minimal alteration of nuclear architecture and may lead to the selection of a small number of meiocytes, but at the same time, does not have a substantial reduce effect on fertility.

### 4.4 Shrew Rb heterozygotes: fertility vs sterility

Complex heterozygotes for multiple Rb translocations are excellent research objects for evaluating the isolating role of these structural chromosome rearrangements. In Rb heterozygotes, meiotic configurations of varied complexity can form during meiosis (see the review by Matveevsky & Kolomiets, 2016), which are expected to be able to influence the progression of meiosis to some extent and to affect reproductive performance of the animals (Searle, 1993; King, 1993 and refs therein). In general, it is postulated that the simple and complex meiotic configurations cause meiotic problems (defects of meiotic chromosomes and meiotic processes) and often result in reduced fecundity of heterozygotes. Lowered fertility of chromosome heterozygotes may be a consequence of increased germ cell death due to meiotic failures and/or formation of aneuploid gametes resulting from nondisjunction of anaphase I and/or embryo inviability. It has turned out that in various species such meiotic configurations give diametrically opposite outcomes, from complete sterility (as, for instance, in some hybrid lemurs, Ratomponirina et al., 1988, or some hybrid mole voles Matveevsky et al., 2020; Kolomiets et al., 2023) to near-normal or full fertility (as in simple heterozygotes in mice Britton-Davidian et al., 1990; Wallace et al., 1992). It should be pointed out that the number and complexity of meiotic configurations may reduce fertility on a spectrum, that is, a greater number of trivalents and more complex multivalents lead to a greater decrease in fertility (a greater increase in the degree of sterility) (lemurs: Dutrillaux & Rumpler, 1977; Ratomponirina et al., 1988; Djlelati et al., 1997; mice: Mahadevaiah et al., 1990, Johannisson, Winking, 1994, Wallace et al., 2002; Nunes et al., 2011). In this regard, common shrew chromosomal races in general and interracial hybrids in particular are some of the best models for research on meiosis progression and fertility in complex heterozygotes.

There have been а number of attempts to evaluate the fertility of heterozygous shrews from different interracial hybrid zones. Often, investigators have focused on such patterns as progression of meiosis, including synapsis and recombination disturbances (see the review by Borodin et al., 2019, and refs therein), chromosome nondisjunction frequencies, and qualitative and quantitative characteristics of spermatogenesis (Garagna et al., 1989a, b; Banaszek et al., 2000a; Jadwiszczak & Banaszek, 2006), and even have analyzed implanted and live embryos (Searle, 1990; Banaszek et al., 2000b).

In the common shrew, many (at least 36) hybrid zones between different chromosomal races are known, and interracial heterozygotes demonstrate diverse meiotic configurations consisting of three to 11 chromosomes (Figure 2). It can be hypothesized that configurations of three chromosomes and those of 8–11 chromosomes make different destructive contributions to meiotic progression and spermatogenesis and in general can have graded effects on fertility. The first supposition that interracial shrew hybrids possess impaired gametogenesis was reported by Aniskin and Volobuev (1980, 1981). The estimated frequency of chromosomal nondisjunctions in hybrids varied widely; in most studies, simple and complex heterozygotes have had 1–13% nondisjunctions (Searle, 1984; Fedyk & Chêtnicki, 2007, 2010; Mercer et al., 1992, Banaszek et al., 2002), and only

Drnholec–Białowieża hybrids have a considerably higher level of nondisjunctions, 27.3% and 40% for CVI and CVII+CIV complex heterozygotes, respectively; but it should be taken into account that few cells (15 and 22 spreads from two males and one male, respectively) have been counted (Jadwiszczak & Banaszek, 2006). No critical problems in the progression of spermatogenesis have been revealed in studies on the ratio of primary spermatocytes (sc) to spermatids (sd) and on the proportion of "defective" tubules in shrew heterozygotes (Garagna et al., 1989b; Narain & Fredga, 1997; Banaszek et al., 2000a; Jadwiszczak & Banaszek, 2006). Despite some differences in spermatogenesis between homozygotes and simple and complex heterozygotes, Garagna et al., (1989b) proposed that they were all fully fertile. Those authors were also mindful that complex heterozygotes could have considerable individual differences, and if we assume that among them, there could be nearly sterile or sterile animals, then this problem could contribute to the features of hybridization between chromosomal races (Garagna et al 1989b). In this regard, of interest are the data showing that if the frequency of CV heterozygotes in a hybrid zone is more than 10%, then reproductive output is low (Banaszek et al., 2000b).

The numbers of spermatozoa have not significantly differed between homozygotes and simple and complex heterozygotes from different hybrid zones (Garagna et al., 1989b; Narain & Fredga, 1997, 1998); accordingly, it has been concluded that all shrews carrying Rb translocations are fully fertile (Garagna et al., 1989b). Our subjective analysis of the number of spermatozoa also did not reveal differences between shrews belonging to the pure Moscow race and Moscow–Seliger F1 hybrids. Moreover, here for the first time we present results of an analysis of some aspects of the morphology and structure of shrew spermatozoa that show no differences between the pure Moscow race and Moscow–Seliger F1 hybrid individuals.

Thus, no researchers have found heterozygous shrews to be sterile animals. In some of the references cited above, authors reported either a partial decrease in fertility or complete fertility. On the other hand, a "partial/decreased in fertility" is a rather vague term because it does not provide clear-cut numerical data on the final fertility of each animal. Additionally, it is not entirely clear whether a partial reduction in fertility implies a reduction by 5%, 20%, or 40%. One way or another, the Moscow– Seliger F1 hybrids investigated here showed some meiotic signs of a slightly decreased fertility, overall consistent with literature data on other interracial hybrid zones of the common shrew. It should be noted that our results of the SC analysis are based on only two hybrid males, and because interindividual differences among heterozygotes have been documented in other papers, a confirmation of our data on a larger number of hybrid shrews is needed.

## 5 CONCLUSIONS

Results of previous studies on various animal models suggest that an open undecavalent can negatively affect the progression of meiosis. Despite the presence of stretched centromeres in the undecavalent and sporadic cases of synaptic irregularities, they nevertheless do not lead to appreciable cell selection. On the contrary, the silencing of centromeric regions of undecavalent asynapsis, the correct recombination between homologous arms of this long meiotic chain (CXI), and the absence of disturbances in diakinesis suggest that the majority of shrew hybrid spermatocytes successfully pass prophase I checkpoints. The number and presence of morphologically normal active spermatozoa in Rb heterozygotes also confirm that meiotic progression is successful at post-prophase I stages as well. We believe that such complex Rb heterozygotes have near-normal fertility, as documented for some other interracial hybrid zones of the common shrew. Possible low stringency of prophase I checkpoints, proper segregation of homologous chromosomes, and the ability of hybrids to form mature germ cells imply rapid evolutionary fixation and circulation of Rb chromosomes within shrew populations, leading to a variety of chromosomal races. The absence of significant abnormalities in meiosis of Moscow-Seliger complex Rb heterozygotes suggests an unlimited gene flow between parental chromosomal races. Note, this is not congruent with the observed stability of the position of this hybrid zone (examined hybrids were caught in 2015 in the central part of the zone which has been discribed by Bulatova with coauthors (2007) 10 years ago), with bimodality of the most of known hybrid zones, as well as with the negative correlation of the width of the known hybrid zones with the range of karyotypic differences between the hybridising races (Bulatova et al., 2011). Explanation of this incongruence requires further studies. In this regard, it seems to us promising to carry out further comprehensive in-depth studies in the Moscow-Seliger hybrid zone.

## ACKNOWLEDGEMENTS

We thank the Genetic Polymorphisms Core Facility of the Vavilov Institute of General Genetics of the Russian Academy of Sciences for the possibility of using microscopes. We are grateful to G.N. Davidovich, A.G. Bogdanov and their colleagues at the Electron Microscopy Laboratory of the Biological Faculty of Moscow State University for their technical assistance. We are grateful to Irina Bakloushinskaya for valuable advice and comments on improving the paper. The English language was corrected and certified by shevchuk-editing.com. This research was supported by the VIGG RAS State Assignment Contracts, Nos 0092- 2022-0002 and A.N. Svertsov IEE RAS State Assignment Contracts, Nos 0089-2021-0007.

## ETHICS STATEMENT

All applicable international, national and institutional guidelines for the care and use of animals were followed. All experiments were approved by the Ethics Committee of the Vavilov Institute of General Genetics of the Russian Academy of Sciences, Russia (order No. 3 of 10 November 2016).

## DATA AVAILABILITY

Additional data presented in Supplementary materials. Other data supporting this study are available from the corresponding authors on reasonable request.

## Supplementary materials

**FIGURE S1.**
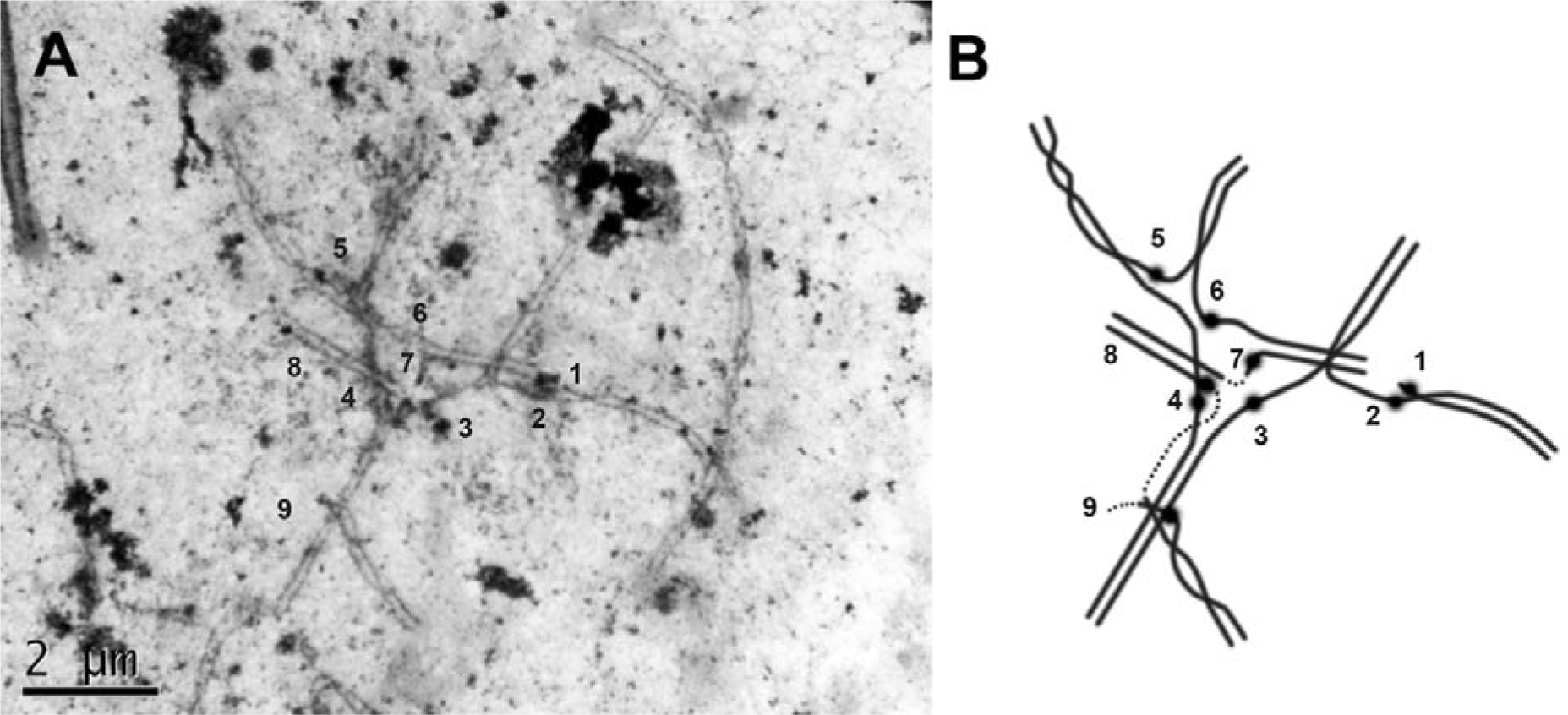
Meiotic configuration in pachytene spermatocytes. A. Electron micrograph; B. Scheme. The numbers correspond to the number of chromosomes in the chain

**FIGURE S2.**
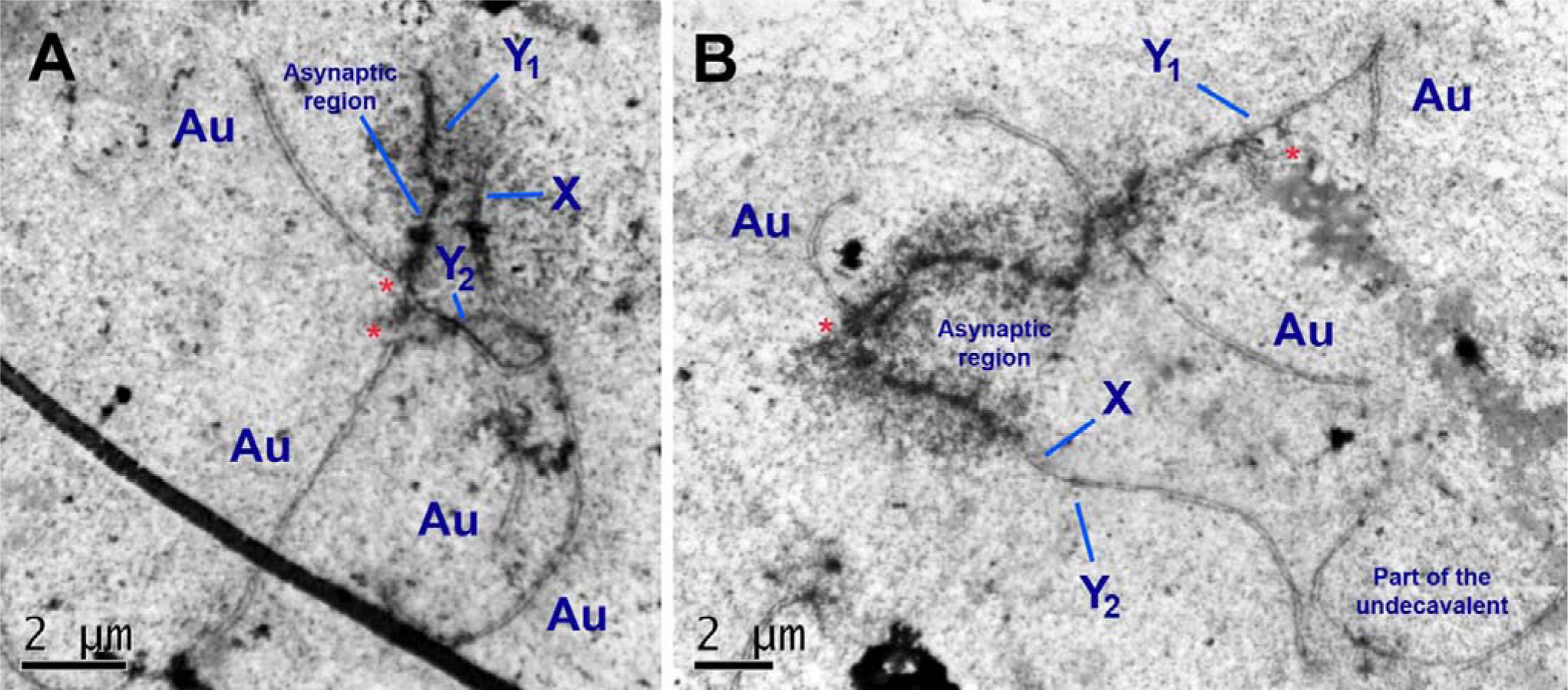
Associations of sex trivalents and autosomes (A, B). Au – autosome. Y_1_, Y_2_, X – chromosomes of sex trivalent. The red asterisk marks the area of contact between the autosome and the asynaptic region of the sex trivalent.

**FIGURE S3.**
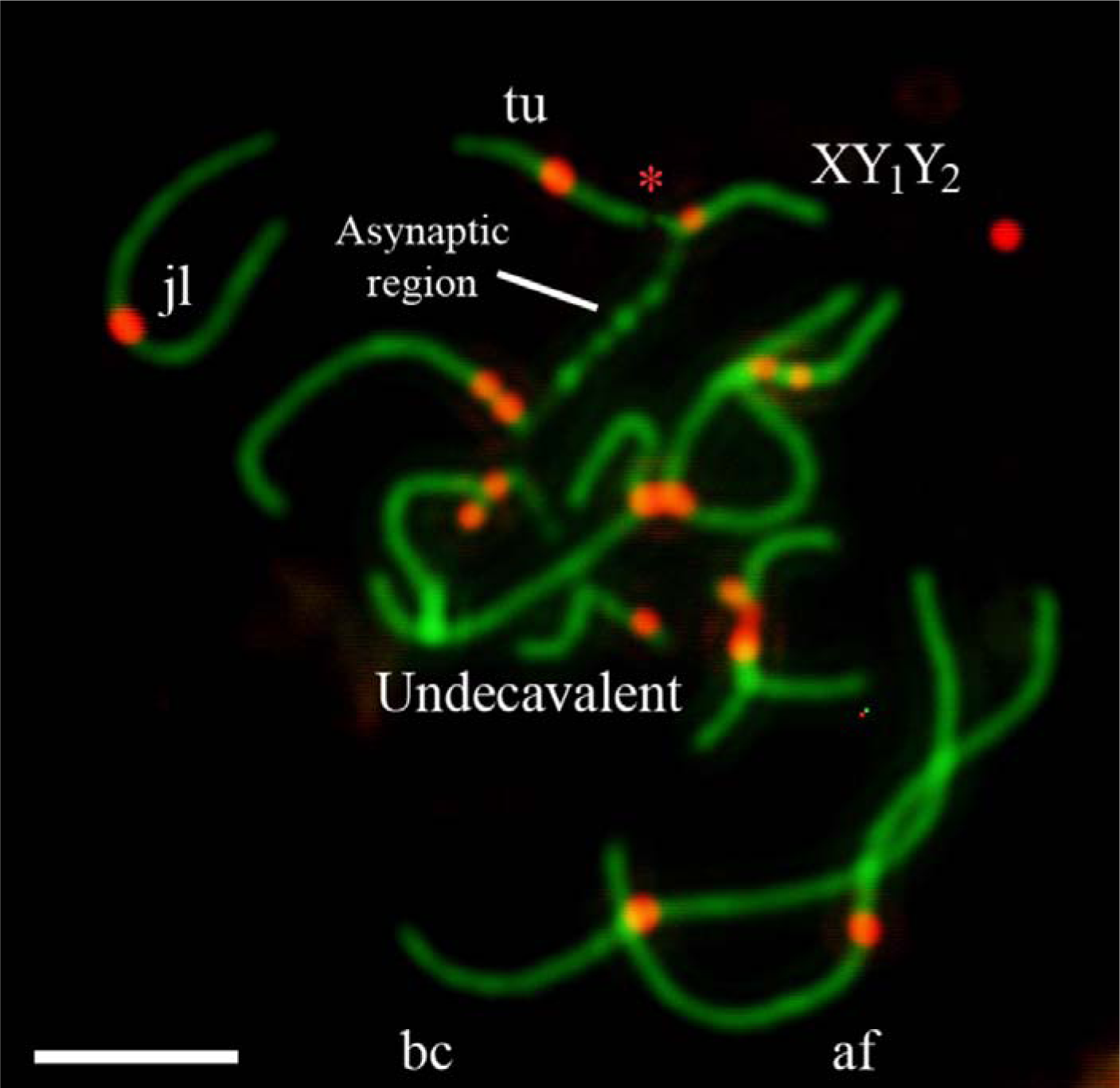
Pachytene spermatocytes of Moscow – Seliger hybrid. The red asterisk marks the area of contact between the tu autosome and the Y1 of the sex trivalent.

**FIGURE S4.**
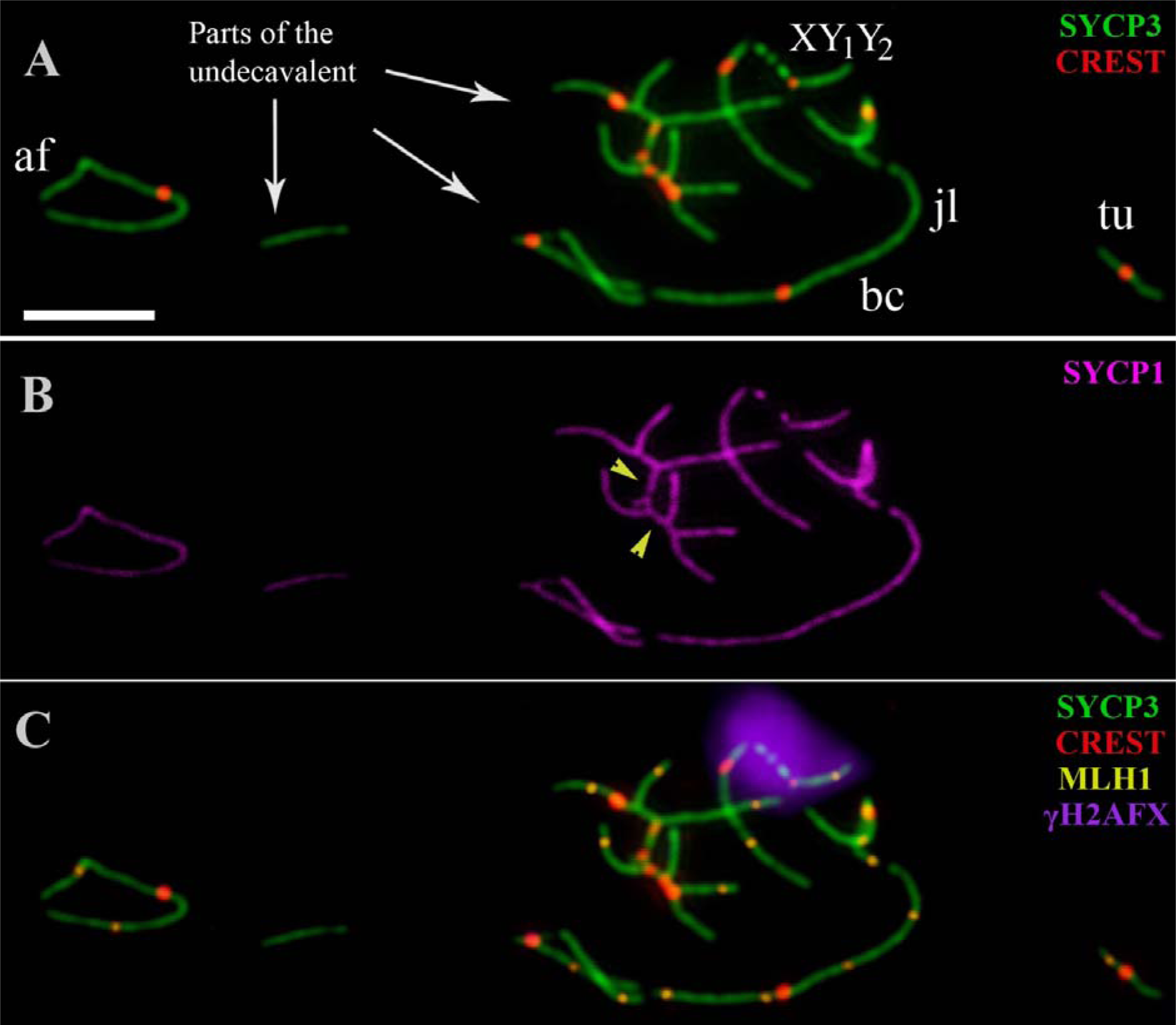
Pachytene spermatocytes of Moscow – Seliger hybrid. SYCP1 (magenta), SYCP3 (green), CREST (red), MLH1 (yellow), γH2AFX (violet) signals are presented. **A.** SYCP3 and CREST signals; **B.** SYCP1 signal; **C.** SYCP3, CREST, MLH1 and γH2AFX (violet) signals. Yellow arrowheads point to the SYCP1 signals in the regions of the heterologous synapsis. Scale bars = 5 μm

**FIGURE S5.**
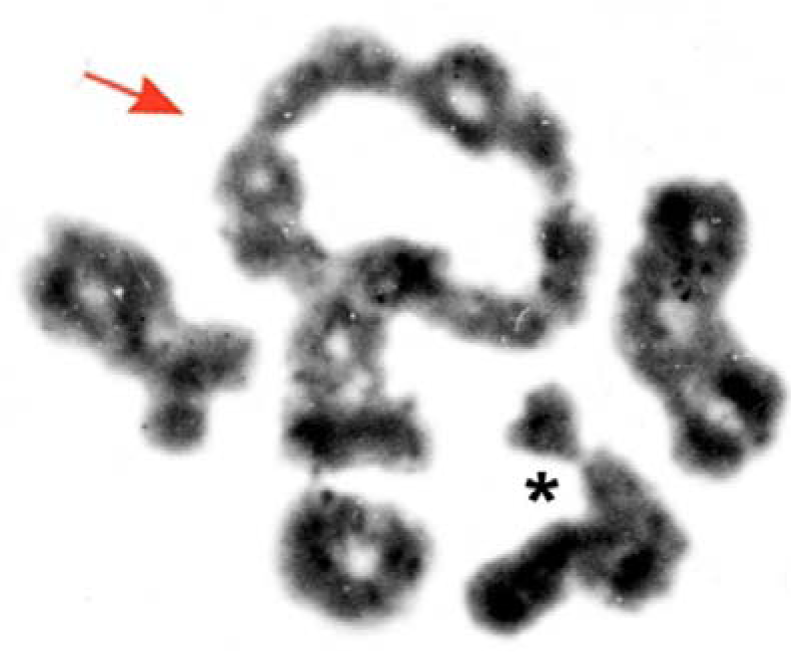
A diakinesis/metaphase I spread of a Moscow–Seliger F1 hybrid male under study (*S. araneus*). The arrow points to a chain-of-eleven configuration. The asterisk indicates a sex trivalent.

**Video S1.**
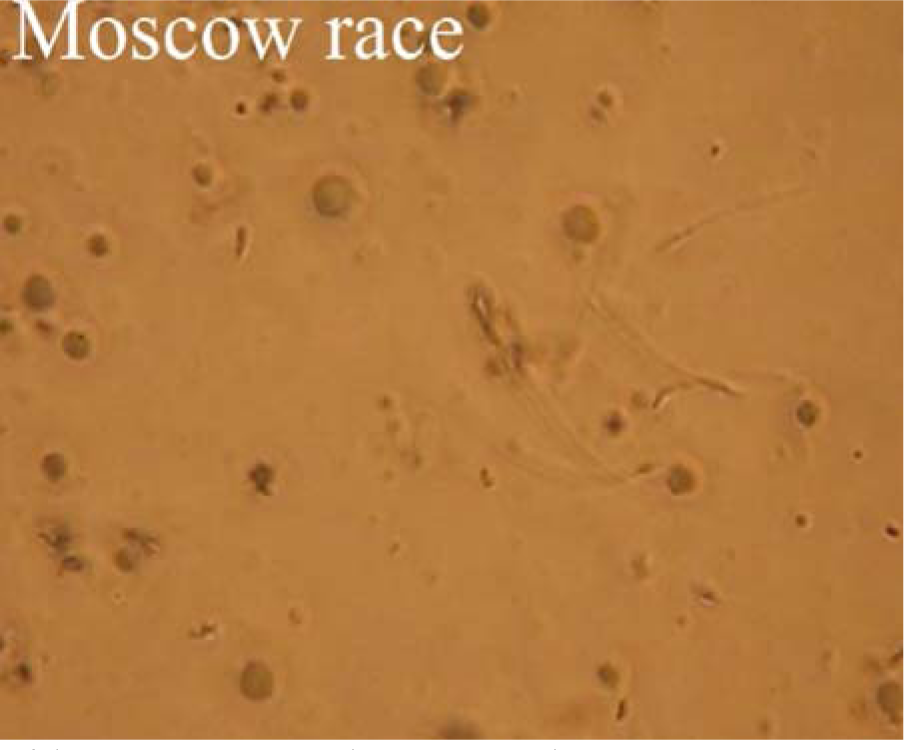
Sperm motility in testicular suspension (Moscow race). Magnificent X100.

**Video S2.**
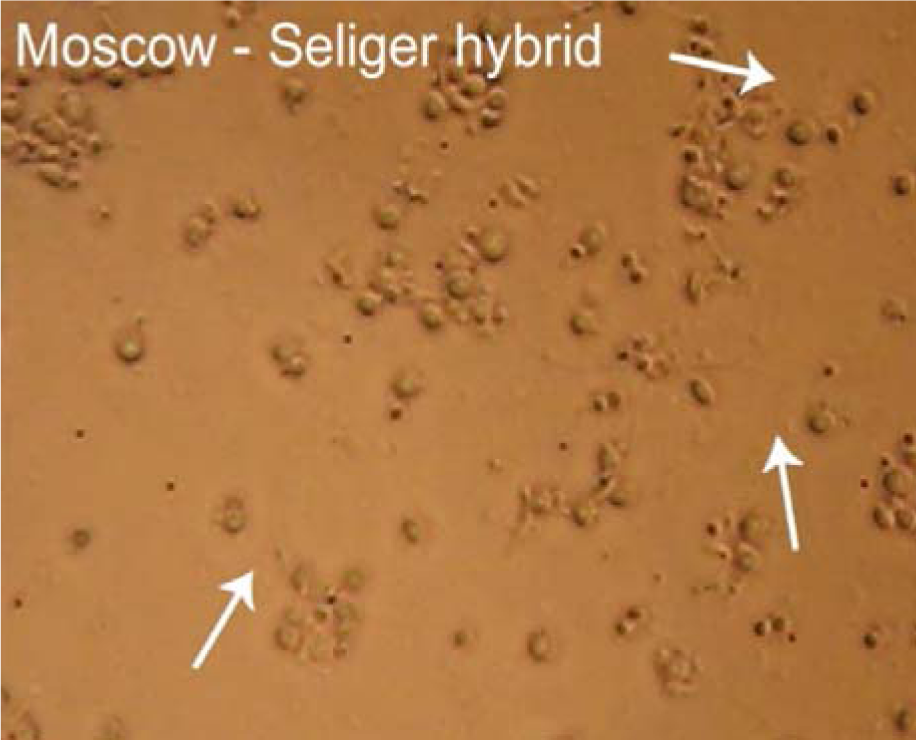
Sperm motility in testicular suspension (Moscow-Seliger hybrid). White arrows point to the active spermatozoa. Magnificent X40.

